# Multi-Layer Functional Metabolomics Reveals Chemical Interactions Between the Plant-Associated Bacterium *Serratia marcescens* and *Phytophthora*

**DOI:** 10.64898/2026.11.17.767058

**Authors:** Daniela Herrera-Rosero, Christian Geibel, Muhammad Waqas, Harald Gross, Silja Mordhorst, Daniel Petras

## Abstract

Oomycete pathogens of the genus *Phytophthora* cause devastating crop losses worldwide, highlighting the need for novel biocontrol agents. Here, we developed a multi-layer functional metabolomics workflow that incorporates an LC–MS/MS-based metal infusion assay in combination with microfractionation-based bioassays against *Phytophthora infestans* and metal-binding assays to map bioactivity from *Serratia marcescens* extracts. MS/MS analysis and feature-based molecular networking supported the annotation of activity-associated metabolites, including prodigiosin, serratiochelins, and an uncharacterized putative serratiochelin analog designated serratiochelin G. Complementary chrome azurol S assays and post-column iron infusion provided evidence of iron-binding properties. Comparative genomic analysis and l-lysine supplementation supported a proposed biosynthetic origin of serratiochelin G involving cadaverine incorporation. This workflow enables metabolite prioritization by integrating non-targeted LC–MS/MS with chromatographically resolved bioactivity and complementary functional assays.

**Table of Contents Graphic:** 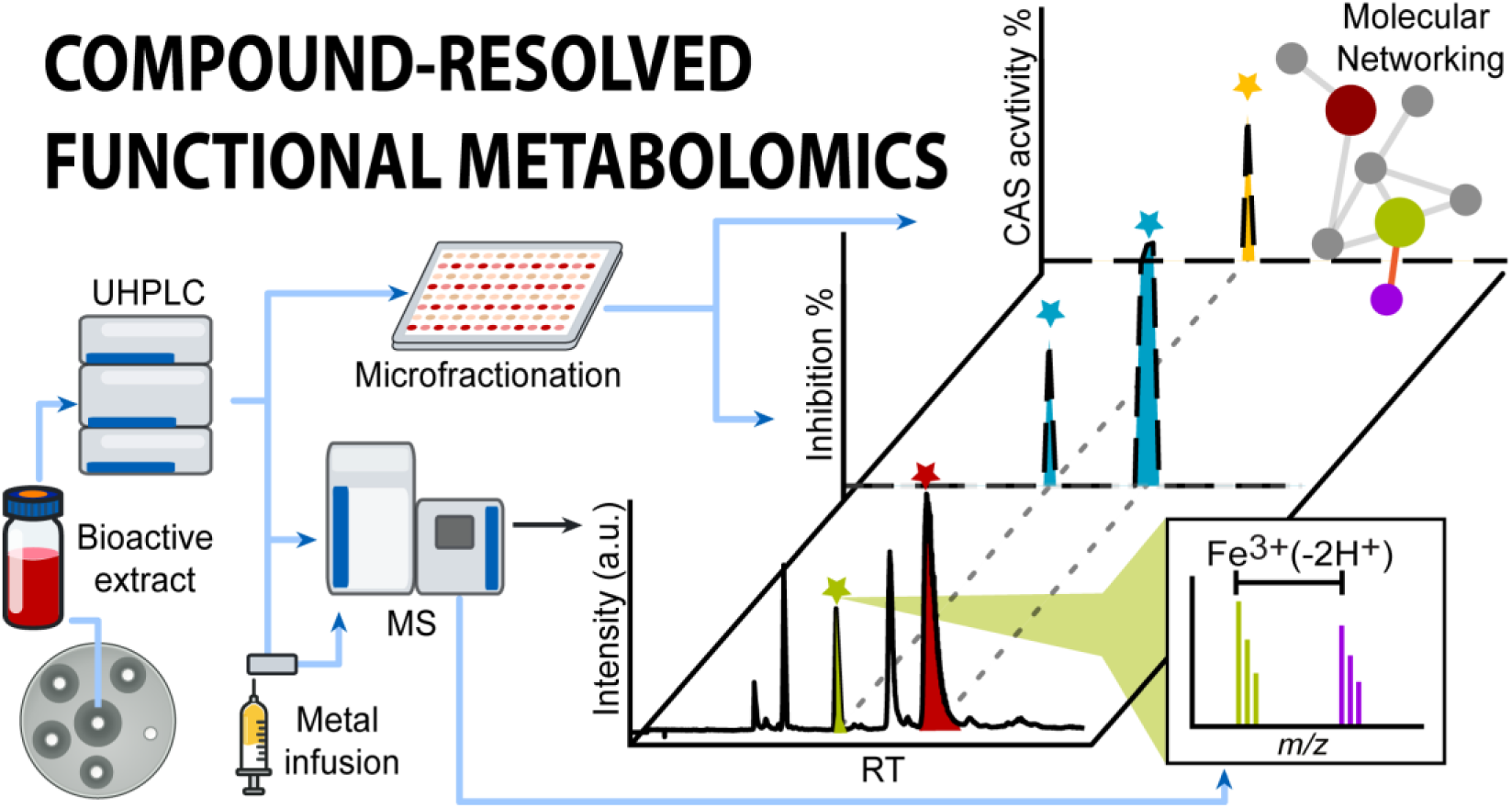

## Introduction

*Phytophthora* species are oomycete plant pathogens associated with destructive diseases across diverse hosts and production systems, representing a major threat to crop production and food security.^1,2^ Within the genus, *Phytophthora infestans* causes late blight of potato and tomato, with large recurring losses in the multi-billion-dollar range annually, underscoring the need for effective control strategies and new intervention concepts.^3,4^

Plant-associated microorganisms can contribute to plant health and protection against biotic stress through the production of specialized metabolites.^5^ Among these microorganisms, members of the genus *Serratia* exhibit diverse beneficial plant-associated functions, including nutrient mobilization, phytohormone production, and activation of plant defenses. Particular strains colonize the rhizosphere and plant tissues, where extracellular enzymes and specialized metabolites can contribute to pathogen suppression. *Serratia marcescens*, a species also recognized as an opportunistic human pathogen,^6^ has, for example, been reported as a plant-associated bacterium with biocontrol activity against a broad range of phytopathogens, including fungal and oomycete pathogens, across different crop systems. Metabolites associated with these activities include prodigiosin, pyrrolnitrin, the anti-oomycete macrolide oocydin A, and biosurfactants such as serrawettins. In addition to producing antimicrobial metabolites, plant-associated *Serratia* can influence nutrient availability through siderophore secretion, linking its extracellular chemistry to competition for iron.^7–9^

Siderophores facilitate microbial iron acquisition by chelating ferric iron, whose limited solubility can restrict its bioavailability.^10^ In *Serratia*, these metabolites include the bis-catecholate serratiochelins and structurally related siderophores assembled through variation in their polyamine backbones.^11^ This chemical diversity motivates the complementary assessment of growth inhibition and iron chelation in *Serratia* extracts. Resolving these functional properties within complex mixtures requires analytical approaches that connect assay responses with chromatographic features and support the prioritization of individual metabolites for further characterization.

Bioassay-guided fractionation typically relies on iterative cycles of bioassay and chromatographic purification, which can require multiple rounds of separation over extended instrument time and substantial manual effort. These workflows are reported to suffer from frequent rediscovery of known compounds after a laborious process, alongside high solvent use and large starting material requirements in some settings.^12^ A further recurring problem is that extracts can yield strong initial bioassay signals while the active compounds are not ultimately isolated during subsequent purification steps.^13^ Co-elution of multiple metabolites can complicate the attribution of bioactivity to individual compounds. As a mitigation, chemical dereplication approaches based on HR-MS, MS/MS, and UV/Vis database querying have been widely used to reduce rediscovery and to improve prioritization, but they do not inherently provide chromatographically resolved alignments between a biological readout and individual LC features.^14^

LC–MS/MS-coupled microfractionation workflows address this alignment problem by collecting many small, time-resolved fractions while acquiring MS data in parallel, enabling correlation between bioassay signals and chromatographic feature profiles.^15^ Ultra-micro-scale fractionation approaches explicitly couple UHPLC–MS separations with microtiter plate bioassays to obtain chemical profile information alongside high-resolution fractionation, with claims of improved chromatographic resolution and reduced fractionation time relative to lower-resolution workflows.^12^ Post-column spotter-based nanofractionation likewise supports high-resolution bioactivity profiling by depositing LC effluent into high-density plate formats and enabling flexible incubation conditions without sacrificing chromatographic resolution due to diffusion during incubation.^16^ A microspotter-based implementation further describes a high-speed fractionation device operating at approximately 1 Hz that spots LC effluent onto a microfluidic paper-analytical device in parallel with LC–MS/MS acquisition, followed by overlay with luminescent bioreporter strains and correlation of luminescence signals to MS chromatograms to localize bioactive components in complex mixtures.^15^

Here, we expanded a high-frequency LC–MS/MS-coupled microspotter workflow^15^ to liquid *Phytophthora* bioassays and extended it with complementary profiling of iron-binding metabolites. Using *S. marcescens* extracts as a model system, chromatographically resolved bioactivity was integrated with chrome azurol S (CAS)-based iron-binding screening, MS/MS molecular networking, and post-column Fe³⁺ infusion to prioritize bioactive and metal-binding features.

### Experimental Section

#### Microorganisms and Maintenance

*Serratia marcescens* DSM 1636 and DSM 30121 were obtained from the German Collection of Microorganisms and Cell Cultures (DSMZ, Braunschweig, Germany), maintained on LB agar and propagated in LB broth for inoculum preparation. *Phytophthora infestans* strain CRA208m2 was obtained from the culture collection of the University of Tübingen and maintained on V8-juice agar. V8 medium was prepared by combining 200 mL of tomato juice, 3 g of CaCO₃, 0.05 g of β-sitosterol, and 15 g of agar, followed by adjustment to 1 L with Milli-Q water and to pH 7.2 ± 0.2. Clarified V8 (cV8) medium for liquid bioassays was prepared by centrifuging 40 mL of tomato juice (4000 rpm, 30 min), collecting the clarified supernatant, and supplementing it with CaCO₃ (0.6 g) and β-sitosterol (0.01 g). The volume was adjusted to 200 mL with Milli-Q water, and the medium was sterilized by autoclaving.

#### Fermentation, Extraction, and Solid-Phase Fractionation of Extracellular Metabolites

*S. marcescens* DSM 1636 and DSM 30121 were cultivated for metabolite production in M9 minimal medium prepared from 200 mL of a 5× M9 salts stock containing Na₂HPO₄·7H₂O (64 g/L), KH₂PO₄ (15 g/L), NaCl (2.5 g/L), NH₄Cl (5 g/L). The medium was supplemented with MgSO₄ (2 mM), CaCl₂ (0.1 mM), and glucose (0.4% *w*/*v*) and adjusted to a final volume of 1 L with Milli-Q water. Cultures were incubated at 27 °C for 48 h under shaking conditions, and the biomass was removed by centrifugation. For the l-lysine supplementation experiment, cultures were grown under the same conditions with l-lysine concentrations of 0, 5, or 20 mM. The resulting clarified supernatants were extracted following the procedure described below. LC–MS/MS analysis of these samples is detailed in the Supporting Information. The clarified supernatants were extracted with ethyl acetate at a supernatant-to-ethyl acetate ratio of 1:2 (*v*/*v*). The mixtures were shaken thoroughly, allowed to separate, and the organic phase was collected. Solvent was removed under reduced pressure using a rotary evaporator to obtain crude extracellular extracts. Between 0.4 and 0.5 g of crude extract were fully resuspended in 20% methanol and subjected to solid-phase extraction (SPE) on Strata™-XL polymeric reversed-phase columns (Phenomenex, 10 g sorbent/60 mL reservoir, 100 μm particle size). Columns were preconditioned with 120 mL of methanol and equilibrated with the same volume of 20% methanol before sample loading. Samples were eluted sequentially with 120 mL of each methanol–water solution, with methanol contents ranging from 20% to 100%. The resulting fractions were evaporated under reduced pressure at 40 °C and stored at −20 °C until further use.

#### Activity Assays

SPE fractions from *S. marcescens* and purified serratiochelins A and B were tested at concentrations of 100, 50, 25, 12.5, 6.25, 3.125, and 1.5625 µg/mL. Standard prodigiosin (AdipoGen Life Sciences) was tested at concentrations of 100, 37.500, 14.063, 5.273, 1.978, 0.742, 0.278, 0.104, 0.039, 0.015, and 0.005 µg/mL. Test solutions were prepared in cV8 medium, with a final DMSO concentration of 0.1% (*v*/*v*) in the assay wells. Each well of a 96-well plate was loaded with 100 µL of the corresponding test solution and 50 µL of a *P. infestans* sporangia suspension (2 × 10⁴ sporangia/mL). All reported compound concentrations refer to the final assay volume of 150 µL after addition of the sporangial suspension. DMSO controls contained 0.1% DMSO in cV8. Plates were incubated at 17 °C for six days, and growth was quantified by measuring OD₆₀₀. Growth inhibition percentages were calculated relative to the mean untreated growth control as 100 × (1 − *OD_sample_*/*O<u>D</u>_control_*). Concentration–response data from three replicates were fitted to a four-parameter logistic (4PL) model by nonlinear least-squares minimization using MATLAB R2025b (MathWorks). IC₅₀ values were calculated by solving the fitted model for the concentration corresponding to 50% growth inhibition relative to the mean growth control, following the definition of absolute IC₅₀ described by Sebaugh,^17^ and were reported only when within the tested concentration range.

#### Microspotter Hardware Setup

The microspotter used in this study followed the general design described by Geibel *et al*.,^15^ consisting of a computer-controlled three-axis positioning platform (STEPCRAFT D.600) equipped with a capillary-based dispensing system. A PEEK capillary (0.13 mm i.d.) was mounted on the positioning arm to enable accurate delivery of the LC eluent. For this work, the system was configured to deposit fractions directly into 96-well microplates, providing a high-frequency microfractionation format compatible with subsequent liquid bioassays. The microspotter traversed the 96-well plate using a serpentine (“snake”) pattern, alternating the direction of movement between successive rows (Figure S1).

#### LC–MS/MS and Microspotter Instrumental Setup

Chromatographic separations were performed on a Kinetex® EVO C18 column (150 × 4.6 mm, 2.6 µm, 100 Å; Phenomenex) using a binary solvent system consisting of water + 0.1% formic acid (*v*/*v*) as mobile phase A and acetonitrile + 0.1% formic acid (*v*/*v*) as mobile phase B. The flow rate was 1.0 mL/min, and the gradient was programmed as follows: 0.00 min, 5% B; 1.00 min, 35% B; 3.00 min, 40% B; 5.00 min, 42% B; 7.75 min, 90% B; 10.00 min, 99% B; 13.00 min, 99% B; 13.10 min, 5% B; held at 5% B until 16.00 min. Post-column, the eluent passed through a time-programmable 6-port valve that directed the flow either to waste or to a T-piece splitting the stream between the mass spectrometer (20%) and the microspotter line (80%). The microspotter inlet was connected to a second 6-port valve controlling delivery to the spotting head. During microfractionation, the 80% microspotter flow was deposited continuously into 96-well microplates over the first 630 s of the chromatographic run, synchronized with LC acquisition. The mass spectrometer (Q Exactive HF, Thermo Fisher Scientific) was operated in data-dependent acquisition (DDA) mode with positive electrospray ionization (HESI), acquiring full-scan MS spectra over *m*/*z* 150–2000 followed by MS/MS of selected precursor ions. Source parameters were as follows: spray voltage 3.5 kV, capillary temperature 250 °C, probe heater temperature 400 °C, sheath gas 50 AU, auxiliary gas 12 AU, and S-Lens RF level 50.

#### Plate-Based Bioassays of Fractionated Samples

Following microfractionation, 96-well plates were dried in a SpeedVac concentrator and subsequently used for liquid bioassays. Each well was supplemented with 100 µL of cV8 medium containing ampicillin (250 mg/L), rifampicin (10 mg/L), and nystatin (19 mg/L) and inoculated with 50 µL of a *P. infestans* sporangia suspension (2 × 10⁴ sporangia/mL). Plates were incubated at 17 °C for six days, and growth was quantified by measuring OD₆₀₀. As validation controls for the microfractionation–bioassay workflow, pyraclostrobin and an 80% methanol solution were processed and spotted under identical conditions and evaluated in parallel. Bioactivity measurements were performed in two independent experiments, each conducted in duplicate.

#### Chromatographic Peak Correlation and Feature-Based Molecular Networking Data Processing

Retention time alignment between microfractionation wells and LC–MS/MS data was performed by extracting the total ion chromatogram (TIC) and MS¹ scan start times using pyOpenMS. A bioactivity chromatogram was generated by calculating growth inhibition for each well as 100 × [1 − (*OD_sample_*/*O<u>D</u>_control_*)], where *O<u>D</u>_control_* represents the mean OD₆₀₀ of the control plate. The resulting inhibition trace was overlaid with the TIC to identify retention-time windows associated with biological activity. Raw LC–MS/MS data were converted to mzML format using ProteoWizard MSConvert and processed in mzmine version 4.5.37^18^ following a standard Feature-Based Molecular Networking (FBMN) workflow, including mass detection, chromatogram building, peak deconvolution, isotope grouping, feature alignment (10 ppm *m*/*z*, 0.15 min RT tolerance), gap filling, and GNPS FBMN export. The resulting feature quantification table and MS/MS spectral summary were uploaded to GNPS2 for FBMN analysis. Library-based spectral matching was performed using GNPS reference libraries (cosine ≥ 0.7; ≥ 5 matched peaks; top 1 hit). Analog search was not enabled in this workflow. Molecular networks were visualized in Cytoscape (version 3.10.3). Full GNPS2 workflow parameters are provided in the Supporting Information.

#### Microspotter–CAS Assay for Detecting Iron-Binding Compounds

Following microfractionation, 96-well plates were dried in a SpeedVac concentrator. A chrome azurol S (CAS) reagent was prepared according to the original formulation.^19^ CAS (0.06 g) was dissolved in 50 mL of deionized water, FeCl₃·6H₂O (0.0027 g) was dissolved in 10 mL of 10 mM HCl, and HDTMA (0.073 g) was dissolved in 40 mL of deionized water. Solutions were combined to generate the blue CAS–Fe–HDTMA complex, which was diluted 1:10 with deionized water immediately before use. Each well was loaded with 50 µL of deionized water followed by 50 µL of the diluted CAS reagent. The color change developed within a few minutes, and absorbance was measured at 600 nm. A decrease in absorbance relative to reagent-only controls indicated Fe³⁺ removal from the CAS complex, consistent with the presence of iron-binding metabolites in the corresponding LC fractions. For chromatographic correlation, CAS activity was calculated relative to a reagent blank containing 50 µL of deionized water and 50 µL of diluted CAS reagent, as %CAS = 100 × [(A₆₀₀,blank − A₆₀₀,sample)/A₆₀₀,blank]. Wells showing ≥30% CAS activity were considered CAS-positive. Retention times were assigned to wells using the same RT–spotting alignment procedure described above (based on MS¹ scan start times extracted with pyOpenMS), generating a CAS activity trace. This %CAS trace was overlaid with the LC–MS TIC to identify retention time windows in which CAS-positive wells coincided with chromatographic peaks.

#### Native Metabolomics with Iron Infusion

LC–MS/MS analyses with post-column iron infusion were performed using an Agilent 1290 Infinity II UHPLC system coupled to a Bruker Impact II QTOF mass spectrometer. Chromatographic separation was carried out on a Phenomenex Kinetex C18 1.7 µm, 50 × 2.1 mm column using mobile phases A (H₂O + 0.1% formic acid) and B (acetonitrile + 0.1% formic acid), following the gradient 0.0 min: 5% B → 10.0 min: 100% B → 12.0 min: 100% B → 13.0 min: 5% B, with 2 min re-equilibration. To enable detection of metal–ligand complexes, the LC eluent was mixed post-column with a FeCl₃ solution (Fe³⁺ as FeCl₃·6H₂O) at 5 mM, following the post-column metal-infusion strategy described by Aron *et al*.^20^ Iron solution was delivered at 0.3 mL/h using a syringe pump through a T-junction placed immediately before the ESI source. This configuration maintains chromatographic separation intact while allowing metal–ligand complex formation immediately before ionization, consistent with the rapid (millisecond) kinetics of metal coordination reported in the original method. LC–MS/MS data were centroided and converted to .mzML before processing in mzmine. Feature detection, chromatogram deconvolution, and alignment were performed in mzmine using a dedicated workflow for Fe-associated ion detection. Ion Identity Molecular Networking (IIMN) was then used to group co-eluting features based on retention time and peak-shape similarity. Ion identity rules included protonated ions and Fe³⁺-associated ions using a theoretical mass offset of Δ*m*/*z* = 52.9115. The mzmine output was networked in GNPS2 using the FBMN workflow with supplemental ion-identity edges for IIMN. Networks were visualized in Cytoscape (version 3.10.3). Detailed IIMN data-processing parameters are provided in the Supporting Information.

#### Genome Mining and Comparative Genomic Analysis

The publicly available genome sequence of *S. marcescens* ATCC 13880, corresponding to DSM 30121, was obtained from ATCC (assembly ID: 9a62beac2315450e; genome ID: 6f11a62a8ec14fc5). Biosynthetic regions were predicted using the antiSMASH web server v8.0.4 with detection strictness set to “relaxed”.^21^

Region 1.1 was exported in the GenBank format and compared with two previously described serratiochelin-associated loci from *Serratia plymuthica* V4 (GenBank accession CP007439.1), encompassing locus tags sch_04710–sch_04735 and sch_19070–sch_19130.^11,22^ Comparisons were performed using clinker through CAGECAT v1.0, with a minimum amino acid sequence identity threshold of 0.3.^23,24^

The same *S. marcescens* DSM 30121 genome assembly was independently annotated using Bakta v1.12.0 with the full v6.0 database to identify candidate lysine decarboxylases.^25^

## Results and Discussion

### Adaptation and Validation of LC–MS/MS-Coupled Microfractionation for Orthogonal Functional Profiling

To identify specialized metabolites that inhibit the growth of *Phytophthora infestans*, we developed a multi-layer functional metabolomics approach that allows the chromatographically resolved profiling of anti-oomycete activity and iron-binding properties. We therefore adapted our recently introduced LC–MS/MS-based high-frequency microspotter platform^15^ to 96-well microplates and combined it with post-column metal infusion experiments.^20^ Post-column LC eluent was split between the mass spectrometer (20%) and the microspotter (80%), and fractions were collected during the first 630 s of the chromatographic run, corresponding to a chromatographic window size of 6.56 s per well. After solvent removal, the collected fractions were either reconstituted in cV8 medium for the liquid *P. infestans* bioassay or treated with deionized water and CAS reagent to assess iron binding, both in the same 96-well format. Technical details of the microspotter configuration are shown in the Supporting Information.

Figure 1 summarizes our analytical workflow and its two complementary layers of functional profiling. Layer 1 corresponds to chromatographically resolved bioactivity profiling using the liquid *P. infestans* assay, whereas Layer 2 provides an orthogonal functional readout of iron-chelating activity using the CAS assay. In both cases, responses from individual wells are mapped back to their corresponding retention-time intervals and integrated with LC–MS/MS features. MS/MS-based molecular networking and feature annotation in GNPS2^26^ provide complementary chemical information for prioritizing features associated with bioactivity, iron binding, or overlapping functional responses. The workflow therefore integrates chromatographic separation, functional profiling, and mass-spectrometric information within a common retention-time framework.

**Figure 1.**
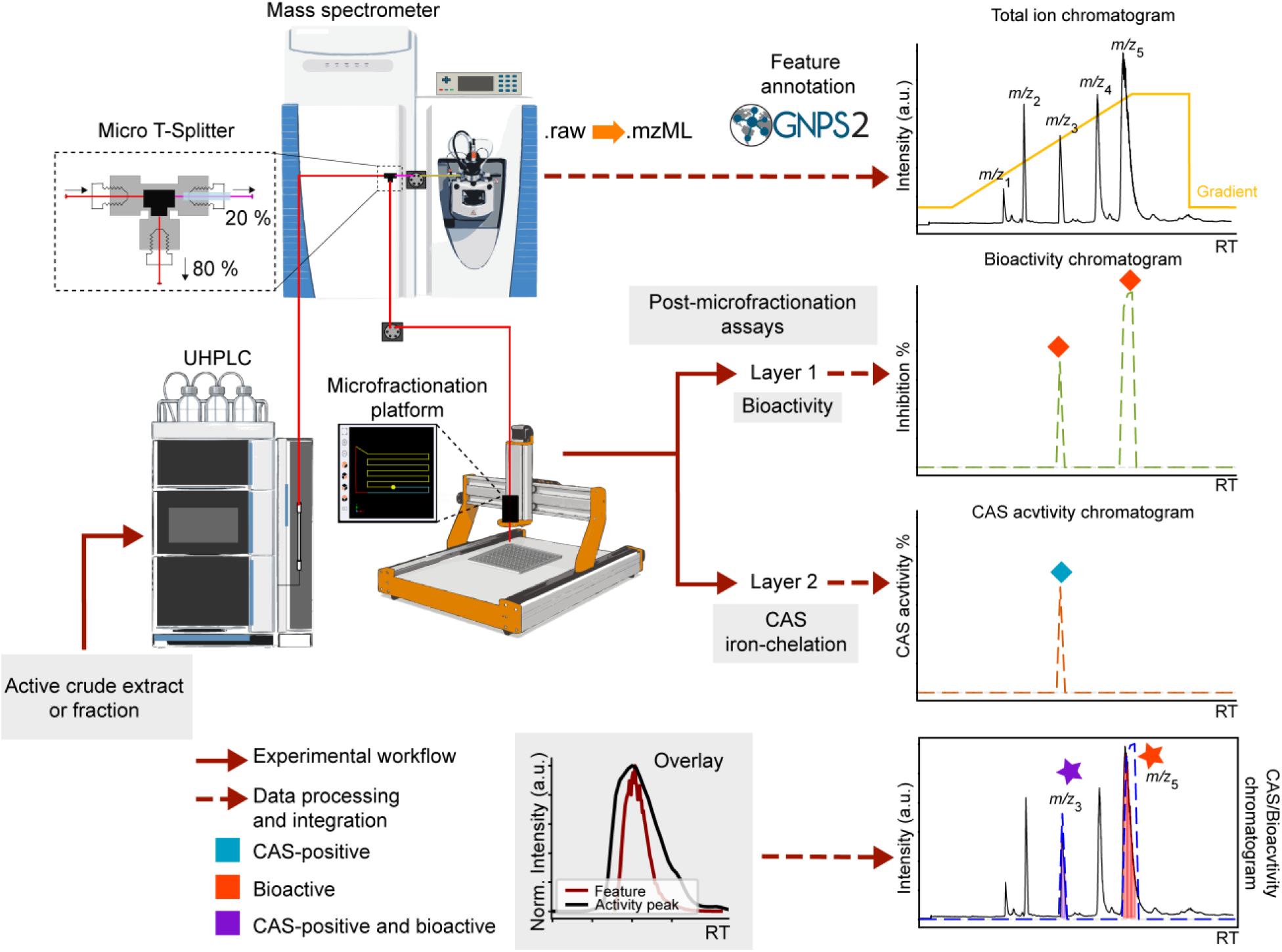
LC–MS/MS-coupled microfractionation workflow for mapping bioactivity and iron-binding metabolites. Extracts or enriched fractions are separated by UHPLC, and the column effluent is split between the mass spectrometer (20%) and the microfractionation platform (80%). Fractions are collected sequentially into 96-well plates, followed by solvent removal and bioactivity or chrome azurol S (CAS) assays. The resulting assay profiles are aligned with chromatographic data to locate bioactive and iron-binding regions. MS/MS-based analysis in GNPS supports metabolite annotation. Solid red arrows indicate experimental steps, whereas dashed red arrows indicate data processing and integration.

Before application to *Serratia* extracts, both functional layers of the workflow were evaluated using appropriate controls. For Layer 1, the broad-spectrum fungicide pyraclostrobin was used as a positive control for the liquid *P. infestans* bioassay, with 80% methanol serving as the solvent control. Pyraclostrobin activity was recovered within a restricted series of consecutive wells corresponding to its chromatographic elution window, whereas the solvent control showed no comparable inhibition signal (Figure S2). These results confirmed that bioactivity could be retained through chromatographic separation and microfractionation and subsequently recovered as a retention-time-resolved inhibition profile in the liquid *P. infestans* assay. For Layer 2, deferoxamine was used as a positive control for the CAS assay to verify that iron-chelating activity could likewise be detected following microfractionation (Figure S3).

### Bioactivity-Resolved Profiling and Prioritization of Serratiochelin-Related Metabolites

Preliminary dual-culture assays showed that *Serratia marcescens* DSM 30121 and DSM 1636 inhibited the growth of *P. infestans* (Figure S4). Dual-culture assays were performed as described in the Supporting Information. A comparison of intracellular and extracellular extracts from DSM 30121 showed visibly greater suppression of *Phytophthora* growth by the extracellular extract (Figure S5), supporting further investigation of metabolites released into the culture supernatant. Culture supernatants from both strains were subsequently extracted and fractionated by reversed-phase solid-phase extraction (SPE) using stepwise methanol elution. The resulting fractions displayed distinct activity profiles against *P. infestans*, with the 80% MeOH SPE fraction showing the highest potency for both strains, with IC₅₀ values of 5.6 μg/mL for DSM 30121 and 2.6 μg/mL for DSM 1636. The corresponding 90% MeOH fractions were also active, with IC₅₀ values of 9.9 and 6.0 μg/mL, respectively, whereas the 70% MeOH fractions showed lower potency (30.8 and 14.1 μg/mL, respectively). Fractions eluted with 20–50% MeOH did not reach 50% inhibition within the tested concentration range, while the 60% and 100% MeOH fractions showed substantially weaker activity. Dose-response curves and calculated IC₅₀ values are provided in Figure S6 and Table S1. Based on these profiles, the 80% MeOH SPE fractions from both strains were prioritized for LC–MS/MS-coupled microfractionation and subsequent testing against *P. infestans*.

For DSM 30121, the resulting microfractions produced spatially distinct inhibitory responses across the assay plate, whereas the corresponding solvent-control plate showed no localized reduction in growth (Figure 2A). Growth inhibition for each well was calculated from OD₆₀₀ measurements relative to the control and assigned to the retention-time interval represented by that well. The resulting inhibition profile was then overlaid with the total ion chromatogram, providing direct temporal alignment between the biological response and the LC–MS data (Figure 2B). A threshold of 50% inhibition was used to prioritize the most pronounced responses. Six discrete bioactivity-associated retention-time windows exceeded this threshold, comprising either individual wells or groups of consecutive wells. The corresponding well positions are indicated directly on the bioactivity chromatogram, while the color-coded outlines in Figure 2A and shaded chromatographic regions in Figure 2B provide a direct visual connection between the plate-based response and its chromatographic position. The agreement between the visibly inhibited wells and the reconstructed inhibition trace supported the use of the microfractionation profile for subsequent feature prioritization. The 80% MeOH SPE fraction from DSM 1636 was analyzed in parallel using the same workflow, and its complete microfractionation profile is provided in the Supporting Information (Figure S7).

**Figure 2.**
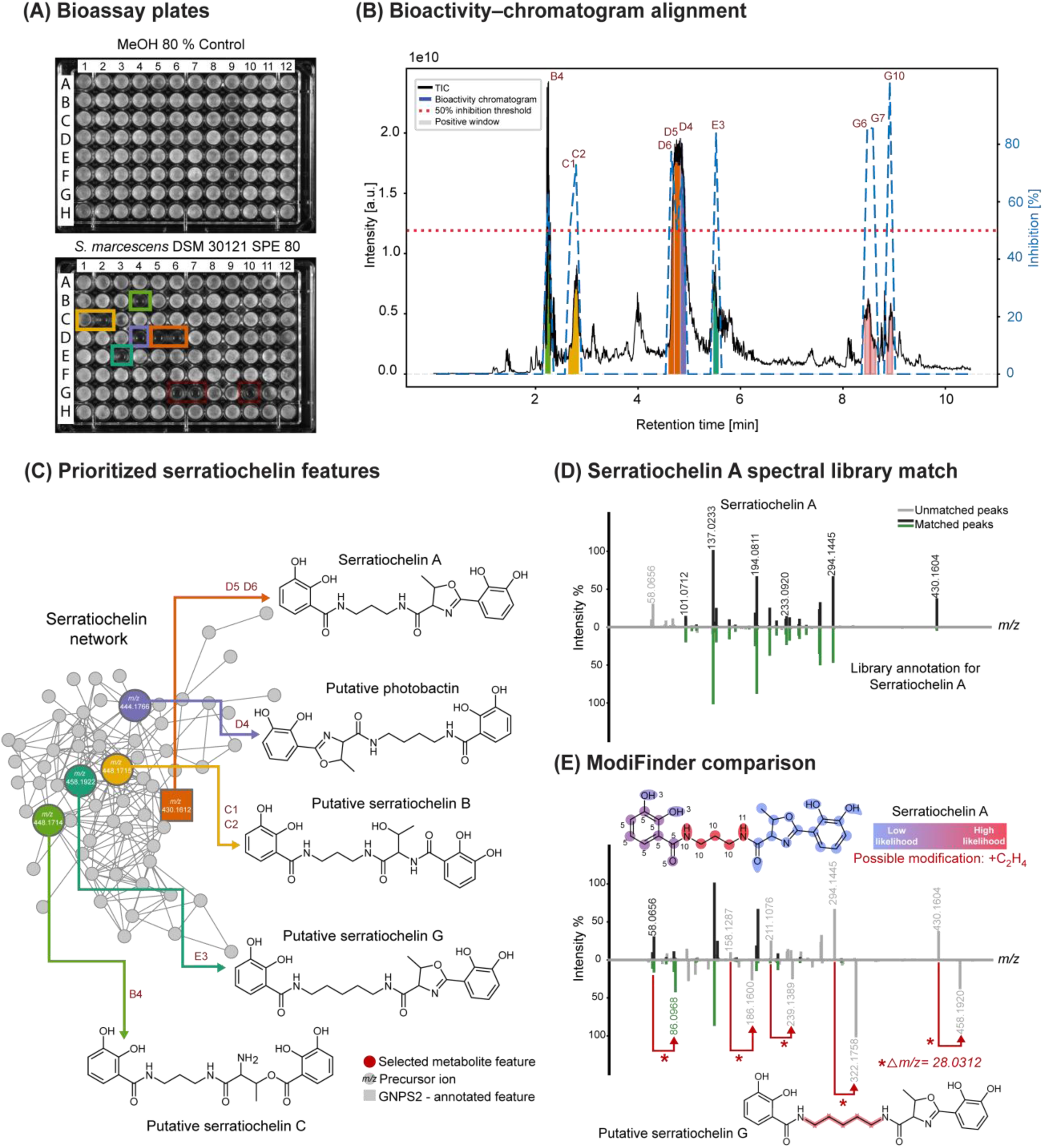
Bioactivity-guided profiling and structural annotation of serratiochelins from *Serratia marcescens* DSM 30121. (A) Microplate bioactivity assay following microfractionation of the 80% MeOH SPE fraction (bottom), with the corresponding solvent control (top). Colored outlines highlight selected active fractions. (B) Total ion chromatogram (TIC) overlaid with the bioactivity profile; shaded regions mark active retention-time windows, labeled with the corresponding well positions. (C) Serratiochelin molecular network obtained by feature-based molecular networking. Nodes represent LC–MS features, and edges indicate MS/MS spectral similarity. Selected nodes are labeled with precursor ion values and linked to their annotated or proposed structures. Colors connect these features with the highlighted fractions in A and B. (D) Mirror plot comparing serratiochelin A (top) with the GNPS2 library spectrum annotated as serratiochelin through Dereplicator+ (bottom). (E) Mirror plot comparing serratiochelin A (top) and serratiochelin G *m*/*z* 458.1922 (bottom), together with the ModiFinder modification-localization map and the proposed serratiochelin G structure. The blue to red scale indicates increasing predicted likelihood of the modification site. Red arrows and an asterisk highlight selected peak pairs separated by Δ*m*/*z* = 28.0312 Da, consistent with a C_2_H_4_ difference. In D and E, matched peaks are shown in black (upper spectra) and green (lower spectra), and unmatched peaks in gray.

Analysis of the LC–MS/MS data from the 80% MeOH SPE fraction of *S. marcescens* DSM 30121 by feature-based molecular networking (FBMN) revealed that the dominant features associated with five of the prioritized bioactive retention-time regions belonged to the same serratiochelin-related molecular family (Figure 2C). FBMN was used because it retains chromatographic feature information, including precursor *m*/*z* and retention time, allowing closely related metabolites to remain represented as distinct nodes within the network.^27^ Five features were prioritized on the basis of their correspondence with the bioactive retention-time windows, their relative abundance within those regions, and their MS/MS relationships: serratiochelin A at *m*/*z* 430.1612, a feature tentatively assigned as photobactin at *m*/*z* 444.1766, and two isomeric features at *m*/*z* 448.1715 assigned as putative serratiochelins B and C based on their distinct chromatographic elution and MS/MS profiles. Putative serratiochelin C was associated with well B4 (RT 2.2–2.3 min), whereas putative serratiochelin B was distributed across wells C1–C2 (RT 2.6–2.8 min). An additional feature at *m*/*z* 458.1922 was designated serratiochelin G. The color-coded connections in Figure 2C link each selected feature to the corresponding bioactive well and retention-time region shown in Figure 2A and B, directly integrating the plate-based inhibition profile with the molecular-network information.

The annotation of serratiochelin A was supported by comparison of its experimental MS/MS spectrum with a GNPS2 library match obtained using Dereplicator+, a computational dereplication approach for natural products,^28^ showing extensive agreement between the fragment-ion patterns (Figure 2D). The assignments of serratiochelins B and C were supported by their precursor masses, chromatographic behavior, and MS/MS relationships within the same molecular family, with the corresponding spectra provided in the Supporting Information (Figure S8). The feature at *m*/*z* 444.1766 was tentatively assigned as photobactin based on its MS/MS relationship and its position within the serratiochelin-associated network. This assignment is biosynthetically plausible because photobactin has previously been reported alongside serratiochelins in *S. plymuthica* V4, where variation in the incorporated polyamine backbone generates structurally related siderophores.^11^ In contrast, serratiochelin G at *m*/*z* 458.1922 did not receive a library annotation but clustered within the same serratiochelin-associated molecular family and corresponded to one of the prioritized bioactive regions. Its structural relationship to serratiochelin A was therefore further examined using ModiFinder.^29^ The MS/MS spectrum of serratiochelin G was compared with that of annotated serratiochelin A, using the serratiochelin A SMILES structure and a 1 ppm mass tolerance. The comparison revealed a precursor mass difference of Δm = 28.0312 Da, consistent with a C₂H₄ difference, while modification-localization analysis assigned the highest likelihood to the central portion of the serratiochelin scaffold (Figure 2E).

Additional later-eluting bioactive regions were also detected in the DSM 30121 microfractionation profile (Figure 2B). These regions contained several LC–MS features and were not associated with a single dominant candidate on the basis of chromatographic alignment and MS/MS information alone. In the parallel analysis of DSM 1636, prodigiosin coincided with a pronounced bioactive region (Figure S7), whereas prodigiosin was not detected in DSM 30121 under the cultivation conditions used here. The persistence of strong bioactivity in DSM 30121 therefore indicates that prodigiosin is not required for the bioactivity observed in DSM 30121 and supports the involvement of additional extracellular metabolites.

Prodigiosin was purified from the 80% MeOH SPE fraction of *S. marcescens* DSM 1636 and its identity was confirmed by comparison with a commercial prodigiosin standard. Serratiochelin A and B were likewise purified and evaluated individually against *P. infestans*. Prodigiosin showed the highest potency, with an IC₅₀ of 0.5 µg/mL, followed by serratiochelin A (IC₅₀ = 7.4 µg/mL) and serratiochelin B (IC₅₀ = 18.5 µg/mL). The corresponding concentration–response curves are provided in Figure S9.

### CAS-Resolved Microfractionation for Iron-Binding Profiling

To complement the bioactivity-resolved profiling and investigate the iron-binding properties of the prioritized chromatographic regions, the microfractionation workflow was coupled to a chrome azurol S (CAS) assay. Analysis of the 80% MeOH SPE fraction from *S. marcescens* DSM 30121 revealed several CAS-positive wells, which were reconstructed as retention-time-resolved CAS response profiles and aligned with the corresponding LC–MS/MS data (Figure 3A,B). Four principal CAS-positive chromatographic regions were observed, all of which corresponded to regions containing members of the serratiochelin-related molecular family identified by MS/MS networking. Particularly pronounced CAS responses were observed in the chromatographic regions containing serratiochelin A and the putative photobactin feature. This observation is consistent with previous studies of *Serratia*, in which CAS-based siderophore activity was associated with serratiochelin production. In *S. marcescens* UMH9, disruption of the *sch* locus resulted in loss of detectable iron-chelating activity in a liquid CAS assay, while subsequent MS analysis identified serratiochelin A as the siderophore produced by this pathway.^30^ Likewise, CAS-positive siderophore production was reported in *Serratia* sp. V4, from which serratiochelins were subsequently structurally characterized.^22^

**Figure 3.**
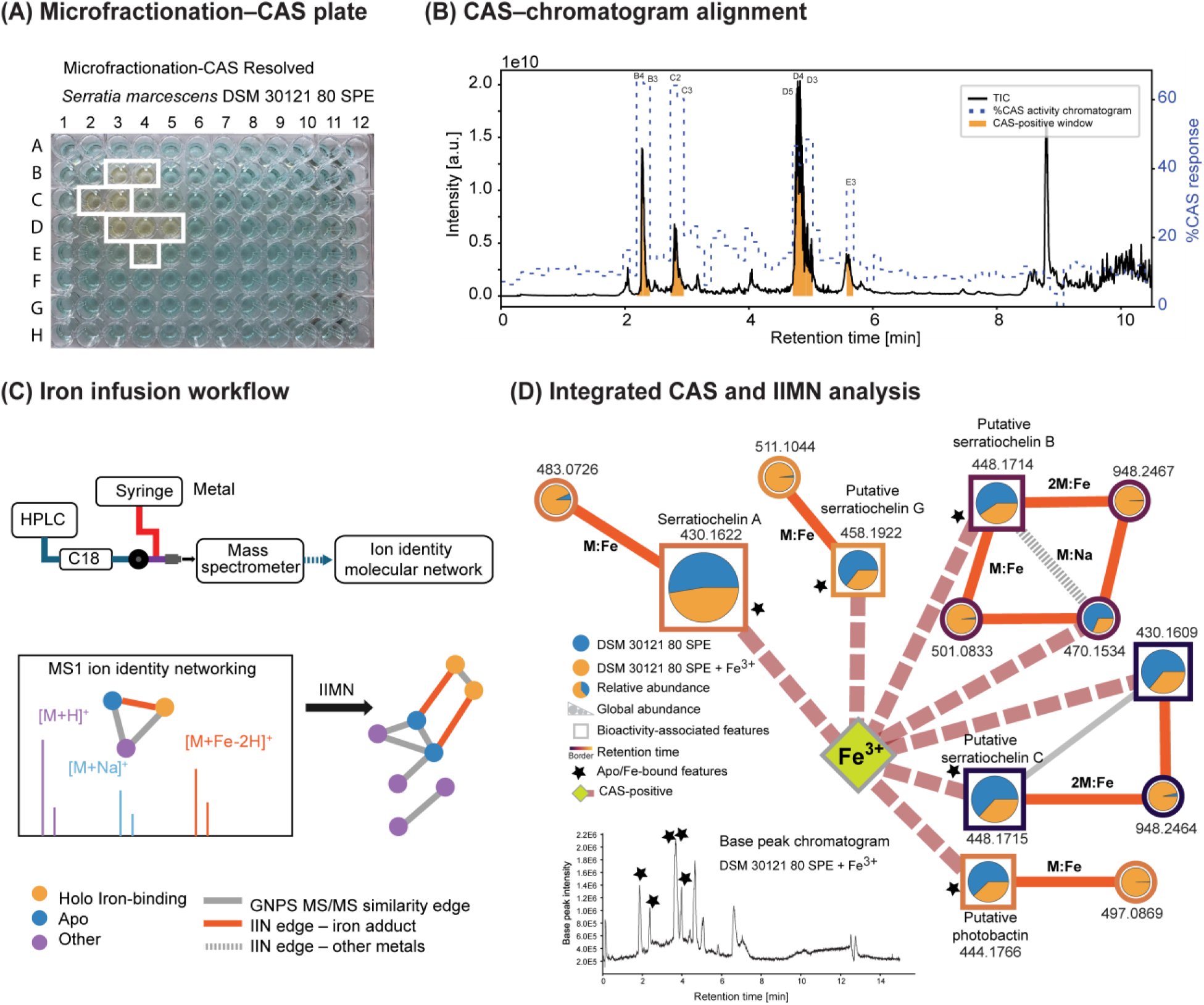
Complementary profiling of iron-binding metabolites by microfractionation–CAS analysis (A, B) and post-column iron infusion (C, D). (A) Chrome azurol S (CAS) assay of the 80% MeOH SPE fraction from *S. marcescens* DSM 30121 following collection with the microfractionation platform. Yellow wells indicate positive CAS responses and are outlined in white. (B) Total ion chromatogram (TIC) overlaid with the CAS response profile; orange shading indicates CAS-positive retention time windows. (C) Schematic of post-column metal infusion and Ion Identity Molecular Networking (IIMN), illustrating the association of co-eluting ion forms through their adduct relationships. (D) Selected subnetworks obtained from the iron-infusion injection showing annotated metabolites and their putative metal complexes, integrating CAS-positive features and Fe-adduct relationships. Nodes are labeled with observed *m*/*z* values. Node size represents global abundance, pie charts show relative abundance in LC–MS injections without (blue) and with (orange) post-column Fe³⁺ infusion, and node borders encode retention time. Labels M and 2M denote proposed 1:1 metabolite-iron and 2:1 metabolite-iron stoichiometries. Stars indicate Fe-adduct-associated features, while the green diamond represents the CAS-positive feature. The inset shows the base peak chromatogram of the Fe³⁺-infused sample, with stars marking the corresponding Fe-adduct-associated retention regions.

In contrast to bulk culture-supernatant CAS measurements, coupling the assay to chromatographic microfractionation provided retention-time-resolved screening of iron-chelating activity. This enabled CAS-positive windows to be directly related to LC–MS/MS features and used as an additional criterion for prioritizing metabolites for subsequent analysis. The magnitude of the CAS response varied across adjacent fractions, consistent with differences in the abundance and/or iron-chelating capacity of metabolites eluting within each chromatographic region. Thus, microfractionation–CAS profiling served as an orthogonal functional screening step that narrowed the regions of interest and guided subsequent feature-level investigation by post-column Fe³⁺ infusion.

The corresponding CAS profiling of the DSM 1636 80% MeOH SPE fraction showed a different response pattern (Figure S10). MS/MS analysis indicated the presence of serratiochelin-related metabolites, including a putative serratiochelin C feature, although these features were detected at lower relative abundance than in DSM 30121 under the conditions analyzed. Notably, the chromatographic region associated with prodigiosin, which coincided with pronounced anti-*Phytophthora* activity in DSM 1636, did not show a corresponding CAS-positive response. Although prodigiosin has been reported to coordinate transition metals such as Cu(II) and Zn(II) through its tripyrrolic scaffold,^31^ this behavior did not translate into detectable Fe³⁺-chelating activity under the CAS conditions used here. The DSM 1636 profile therefore further distinguished bioactivity from iron-chelating activity and reinforced the value of the CAS readout as a complementary functional dimension within the workflow.

The distinction between growth inhibition and iron chelation is relevant because siderophore-mediated effects depend on the biological context. In a maize–peanut intercropping system, *Pseudomonas*-derived pyoverdine improved peanut iron nutrition.^32^ In tomato rhizosphere communities, siderophore-mediated interactions with *Ralstonia solanacearum* ranged from suppression to facilitation, consistent with differences in the pathogen’s ability to use siderophore-bound iron.^33^ Experiments with *Pseudomonas* consortia further linked siderophore-mediated interactions to protection against bacterial wilt, with pronounced pathogen inhibition under iron limitation.^34^ These findings provide a biological rationale for assessing iron chelation alongside growth inhibition. In the present study, however, the overlap between CAS-positive and bioactive chromatographic regions does not establish iron deprivation as the mechanism of *P. infestans* inhibition. Instead, the complementary readouts prioritize metabolites displaying one or both functional properties for further investigation.

### Post-column Iron Infusion Reveals Fe-Binding Serratiochelin-Related Metabolites

Following the retention-time-resolved screening of iron-chelating activity by the CAS assay, post-column Fe³⁺ infusion coupled to electrospray ionization (ESI) mass spectrometry was used to obtain feature-level evidence of metal binding in the DSM 30121 80% MeOH SPE fraction (Figure 3C). This approach builds on native metabolomics strategies developed for the detection of metal-binding metabolites directly in complex biological extracts through the formation of co-eluting metal-associated ion species with characteristic mass relationships to their corresponding metal-free features.^20^ The Fe³⁺-infused and non-infused LC–MS datasets were subsequently integrated using IIMN, which connects co-eluting ion species related by defined adduct and ion-identity relationships within the GNPS molecular networking environment.^35^ In this way, the CAS-based chromatographic screening was complemented by direct MS-based interrogation of individual candidate metal-binding metabolites.

Inspection of the selected IIMN subnetwork revealed Fe-associated ion identities for several members of the serratiochelin-related molecular family (Figure 3D). Serratiochelin A at *m*/*z* 430.1622 ([M+H]⁺) was connected to an Fe-associated feature at *m*/*z* 483.0726, consistent with formation of the [M+Fe−2H]⁺ ion (observed Δ*m*/*z* = 52.9104; expected Δ*m*/*z* = 52.9115). This assignment is also consistent with the previously reported theoretical *m*/*z* value of 483.0729 for ferri-serratiochelin A, with a mass error of −0.62 ppm.^36^ Comparable relationships were observed for the putative serratiochelin G, from *m*/*z* 458.1922 to 511.1044 (Δ*m*/*z* = 52.9122), putative serratiochelin B at RT 2.4 min, from *m*/*z* 448.1714 to 501.0833 (Δ*m*/*z* = 52.9119), and putative photobactin, from *m*/*z* 444.1766 to 497.0869 (Δ*m*/*z* = 52.9103). In each case, the Fe-associated feature was detected predominantly or exclusively following post-column Fe³⁺ infusion, whereas the corresponding metal-free feature remained detectable in both the non-infused and Fe-infused injections. This coexistence of apo and Fe-bound ions after metal infusion is consistent with the behavior reported during development of the native metabolomics workflow, where both forms of a siderophore were observed simultaneously following post-column Fe³⁺ addition.^20^ The node pie charts in Figure 3D therefore visualize the distribution of each LC–MS feature between the two experimental conditions, while node size represents its global abundance.

Additional ion identities further supported the assignment of the serratiochelin-related features and revealed differences in their observed Fe-binding behavior. Putative serratiochelin B at *m*/*z* 448.1714 was also associated with a sodium-containing ion at *m*/*z* 470.1534 (Δ*m*/*z* = 21.9820), consistent with the expected Na⁺/H⁺ adduct relationship, and formed both a monomeric M:Fe complex at *m*/*z* 501.0833 (mass error −0.4 ppm) and a higher-mass feature at *m*/*z* 948.2467 consistent with a 2:1 metabolite–Fe complex ([2M+Fe−2H]⁺). Putative serratiochelin C at *m*/*z* 448.1715 at RT 1.9 min showed a related feature at *m*/*z* 430.1609, corresponding to a loss of 18.0106 Da. Notably, a water-loss ion at *m*/*z* 430.1610 has previously been reported specifically for apo-serratiochelin C in positive-ion ESI,^36^ whereas the corresponding ferri form retained the intact serratiochelin C mass. This behavior is also chemically related to the established interconversion within the serratiochelin family: serratiochelin A was originally characterized as the dehydrated 2-methyloxazoline analog of serratiochelins B/C, and acid-mediated hydrolysis of serratiochelin A produces serratiochelin C.^22^ Under the present post-column infusion conditions, no monomeric M:Fe relationship was detected for putative serratiochelin C; instead, it was associated with a 2M:Fe feature at *m*/*z* 948.2464. Previous studies have nevertheless directly detected the 1:1 ferri-serratiochelin C species at approximately *m*/*z* 501.0844, indicating that the absence of a corresponding monomeric IIMN relationship here should not be interpreted as an inability of serratiochelin C to form a 1:1 Fe complex.^36^

The occurrence of alternative Fe-associated stoichiometries is chemically plausible in the context of the unusual coordination behavior reported for this siderophore family. Serratiochelins are bis-catecholate siderophores, yet serratiochelin A was shown to form a 1:1 Fe complex with predominantly tetradentate coordination involving one catechol, the oxazoline nitrogen, and one hydroxyl group of the second catechol, while a hexacoordinate bis-aqua form was also observed. Moreover, serratiochelin B has previously been reported to exhibit lower affinity for Fe and Ga than serratiochelin A, indicating that structural differences within the family can influence metal-binding behavior.^22^ The 2M:Fe features observed here may therefore reflect alternative Fe-associated assemblies favored under the post-column infusion and native electrospray conditions. Fe-bound 2:1 complexes have also been observed for other chelating metabolites using post-LC metal-infusion metabolomics, demonstrating that such higher-order ion identities can be captured by this analytical strategy.^20^

### Biosynthetic Context and l-lysine Feeding Support a Putative Cadaverine-Derived Serratiochelin Analog

Building on the structural relationship between serratiochelin G and serratiochelin A established above, we next examined whether the proposed cadaverine-derived analog was compatible with the biosynthetic capacity of *S. marcescens* DSM 30121. The additional C₂H₄ unit in serratiochelin G is consistent with replacement of the propane-1,3-diamine backbone of serratiochelin A by cadaverine, prompting investigation of both the serratiochelin biosynthetic machinery and the capacity of DSM 30121 to generate cadaverine from l-lysine. Comparative genomic analysis and precursor-feeding experiments were therefore used to evaluate whether this structural hypothesis was supported independently at the biosynthetic level.

Genome mining of *S. marcescens* DSM 30121 using antiSMASH^21^ identified region 1.1 as a predicted nonribosomal peptide metallophore biosynthetic region, although with low similarity confidence to characterized reference clusters. Previous work in *S. plymuthica* V4 showed that serratiochelin biosynthesis requires two physically separated loci, P1 and P2, located approximately 3 Mb apart and containing enterobactin- and vibriobactin-related biosynthetic genes. Comparison using clinker^23^ showed that homologs corresponding to both loci are colocalized within the single antiSMASH-predicted region 1.1 of DSM 30121 (Figure 4A), representing a genomic organization distinct from that described for *S. plymuthica* V4 while retaining the principal components associated with serratiochelin biosynthesis.^22^ The conserved region included homologs involved in 2,3-dihydroxybenzoate biosynthesis and activation, nonribosomal assembly, siderophore transport, and the stand-alone amide synthase SchH, which has been implicated in polyamine incorporation during serratiochelin-related biosynthesis.^11^ Because a cadaverine-derived analog would additionally require cellular production of cadaverine, the DSM 30121 genome was independently annotated using Bakta.^25^ This analysis identified both CadA and LdcC, annotated as lysine decarboxylases (EC 4.1.1.18), together with the adjacent CadB lysine/cadaverine antiporter and CadC transcriptional regulator associated with the *cad* system. These genes are located outside the serratiochelin biosynthetic region and provide an independent genomic route for conversion of l-lysine to cadaverine (Figure 4C), consistent with previous reports of CadA-mediated lysine decarboxylation and cadaverine production in *S. marcescens.*^37^

**Figure 4.**
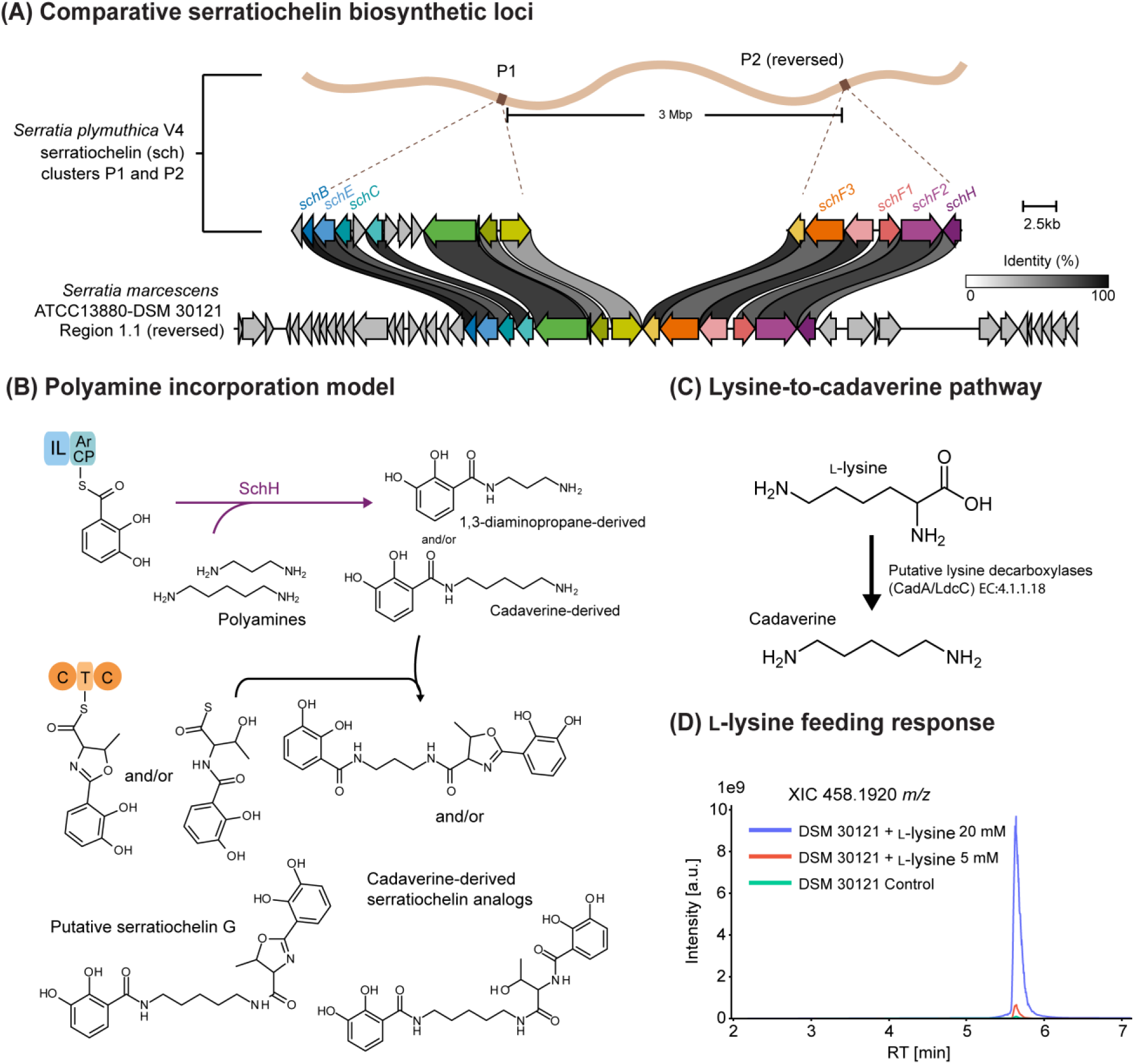
Comparative genomic context and experimental support for a putative cadaverine-derived serratiochelin analog. (A) Comparison of the serratiochelin biosynthetic loci P1 and P2 from *Serratia plymuthica* V4 with antiSMASH-predicted region 1.1 from *Serratia marcescens* DSM 30121, visualized using clinker. Arrows represent coding sequences and their orientation; connecting ribbons indicate amino acid sequence identity between homologous proteins. (B) Proposed biosynthetic scheme for serratiochelin A and cadaverine-derived analogs, illustrating alternative incorporation of propane-1,3-diamine or cadaverine through SchH-mediated acylation and subsequent assembly. (C) l--Lysine decarboxylation to cadaverine, with CadA and LdcC identified as candidate enzymes in the DSM 30121 genome annotation. (D) Extracted ion chromatograms at *m*/*z* 458.1920, corresponding to the putative serratiochelin G, for cultures supplemented with 20 mM, 5 mM l--lysine, or no added l--lysine.

Because cadaverine can be generated through decarboxylation of l-lysine, precursor feeding was used to test whether increasing l-lysine availability influenced the abundance of serratiochelin G. *S. marcescens* DSM 30121 was cultivated with 0, 5, or 20 mM l-lysine, followed by extraction and LC–MS/MS analysis. Extracted ion chromatograms for serratiochelin G at *m*/*z* 458.1920 [M+H]⁺ showed a progressive increase in signal with l-lysine supplementation (Figure 4D). Consistently, mzmine-integrated peak areas (a.u.) increased from 8.17 × 10⁶ in the unsupplemented control to 5.18 × 10⁷ at 5 mM and 1.01 × 10⁹ at 20 mM l-lysine, corresponding to approximately 6.3-fold and 124-fold increases relative to the control, respectively. This pronounced enrichment of the serratiochelin G-associated signal with increasing l-lysine availability is consistent with a biosynthetic relationship between lysine-derived cadaverine and serratiochelin G.

This response is consistent with the genomic identification of the lysine decarboxylases CadA and LdcC in DSM 30121 and with previous observations that *S. marcescens* can produce cadaverine from l-lysine.^38^ Importantly, precursor-directed studies of serratiochelin-related biosynthesis have shown that cadaverine can be accepted by the corresponding polyamine-incorporating machinery, although only partially assembled products were detected and the corresponding full-length analog was not observed.^39^ In contrast, serratiochelin G displays the precursor mass and MS/MS characteristics expected for a complete cadaverine-containing serratiochelin scaffold. The l-lysine-dependent increase observed here therefore provides experimental support for endogenous formation of this putative cadaverine-derived serratiochelin analog in *S. marcescens* DSM 30121.

Molecular networking further revealed additional features associated with the serratiochelin family whose abundance increased following l-lysine supplementation, including the serratiochelin G-related ion at *m*/*z* 476.2031 [M+H]⁺ (Figure S11). These observations suggest that increased precursor availability may influence a broader set of polyamine-containing siderophore-related metabolites rather than serratiochelin G alone.

## Conclusion

This study describes the use of a multi-layer LC–MS/MS-based functional metabolomics workflow for chromatographically resolved profiling of extracellular metabolites from *S. marcescens* against *P. infestans*. Adaptation of the microspotter platform to a liquid oomycete bioassay enabled direct alignment of inhibitory responses with LC–MS/MS features, while complementary microfractionation–CAS profiling and metal infusion MS experiments introduced a second functional dimension related to iron chelation. Application of the workflow to *S. marcescens* DSM 30121 and DSM 1636 revealed distinct activity and metabolite profiles. In DSM 30121, several bioactivity-associated regions were linked to a serratiochelin-related molecular family, whereas prodigiosin represented a prominent activity-associated feature in DSM 1636. Integration of CAS profiling with post-column Fe³⁺ infusion and Ion Identity Molecular Networking provided feature-level evidence for iron binding by serratiochelin A, putative serratiochelin B, photobactin, and the putative serratiochelin analog designated serratiochelin G.

Serratiochelin G emerged as a particularly informative candidate through the combination of multiple independent observations. Its MS/MS relationship to serratiochelin A was consistent with an additional C₂H₄ unit localized to the polyamine-containing region, while native metabolomics supported formation of an Fe-associated ion. Comparative genome analysis identified a serratiochelin-related biosynthetic region in DSM 30121 together with independent lysine decarboxylase pathways capable of generating cadaverine. Consistently, l-lysine supplementation produced a pronounced concentration-dependent increase in the serratiochelin G-associated LC–MS signal. Together, these findings highlight the efficiency of our multi-layer functional metabolomics approach in annotating bioactive metabolites. Ultimately, our data identify serratiochelin G as a putative cadaverine-derived serratiochelin A analog and demonstrate how integration of chromatographic microfractionation, functional assays, molecular networking, metal-infusion metabolomics, and biosynthetic context can strengthen prioritization of bioactive natural products.

## Associated content

### Data Availability Statement

LC–MS/MS datasets generated in this study are available through Zenodo. Workflow-validation datasets are available at https://zenodo.org/records/16637888 and https://zenodo.org/records/13306623. The microfractionation–bioactivity dataset is available at https://zenodo.org/records/15648837, with the corresponding GNPS2 molecular networking results at https://gnps2.org/status?task=219b2652bcc647daaabfc23c4d394ffb. The microfractionation–CAS dataset is available at https://zenodo.org/records/17511886, with the corresponding GNPS2 results at https://gnps2.org/status?task=bf78f8b2208f4557902dacc26fc76a5e. The post-column Fe³⁺-infusion dataset is available at https://zenodo.org/records/23314855, with the corresponding GNPS2/IIMN results at https://gnps2.org/status?task=9cec0027e57b465ea0034a88b4928252 and https://gnps2.org/status?task=4595413309fd439ab924919cc1cc0ca0. The l-lysine feeding dataset is available at https://zenodo.org/records/23334756, with the corresponding GNPS2 molecular networking results at https://gnps2.org/status?task=f7673a9597624be888c24a99787dcd6c. Analysis code and example input files used to generate retention-time-resolved overlays of LC–MS profiles with bioactivity and CAS assay readouts are available at https://github.com/Functional-Metabolomics-Lab/Multi-Layer-Functional-Metabolomics.

## Supporting Information

- Microspotter technical configuration; validation of the microfractionation–bioassay and microfractionation–CAS workflows; dual-culture antagonism and intracellular/extracellular extract comparison; SPE dose–response assays and IC₅₀ values; DSM 1636 microfractionation bioactivity and molecular-networking profiles; supporting MS/MS spectra for serratiochelin-related metabolite annotations; bioactivity validation of prodigiosin and serratiochelins A and B; DSM 1636 microfractionation–CAS profiling; l-lysine-dependent enrichment of serratiochelin-related features; and supplementary methods for dual-culture assays, mzmine processing, GNPS2 FBMN, and IIMN analysis (PDF). G-code for 96-well microspotter fractionation (TXT).

## Acknowledgements

This study was supported by the Deutsche Forschungsgemeinschaft (German Research Foundation, DFG) via the Cluster of Excellence EXC 2124: Controlling Microbes to Fight Infection (CMFI, project ID 390838134). D.H.R. acknowledges funding from the German Academic Exchange Service (DAAD) through the program ‘Research Grants – Doctoral Programmes in Germany, 2023/24’.

## Author Contributions

D.H.R., S.M., and D.P. conceptualized the study. D.H.R., C.G., and M.W. performed the experiments. D.H.R. analyzed the data. H.G., S.M., and D.P. provided materials, instrumentation, resources, and supervision. D.H.R. wrote the manuscript. All authors reviewed, edited, and approved the final manuscript.

## Author Information Corresponding Authors

**Silja Mordhorst** - Cluster of Excellence Controlling Microbes to Fight Infections (CMFI), University of Tübingen, 72076 Tübingen, Germany; Department of Pharmaceutical Biology, University of Tübingen, 72076 Tübingen, Germany; German Center for Infection Research (DZIF), Partner Site Tübingen, 72076 Tübingen, Germany;

**Daniel Petras** - Cluster of Excellence Controlling Microbes to Fight Infections (CMFI), University of Tübingen, 72076 Tübingen, Germany; Department of Biochemistry, University of California Riverside, Riverside, California 72521, United States;

## Authors

**Daniela Herrera-Rosero** - Cluster of Excellence Controlling Microbes to Fight Infections (CMFI), University of Tübingen, 72076 Tübingen, Germany; Pharmaceutical Institute, Department of Pharmaceutical Biology, University of Tübingen, 72076 Tübingen, Germany;

**Christian Geibel** - Cluster of Excellence Controlling Microbes to Fight Infections (CMFI), University of Tübingen, 72076 Tübingen, Germany; Department of Microbial Bioactive Compounds, Interfaculty Institute of Microbiology and Infection Medicine (IMIT), University of Tübingen, 72076 Tübingen, Germany;

**Muhammad Waqas** - Pharmaceutical Institute, Department of Pharmaceutical Biology, University of Tübingen, 72076 Tübingen, Germany;

**Harald Gross** - Cluster of Excellence Controlling Microbes to Fight Infections (CMFI); Pharmaceutical Institute, Department of Pharmaceutical Biology, University of Tübingen, 72076 Tübingen, Germany; German Center for Infection Research (DZIF), Partner Site Tübingen, 72076 Tübingen, Germany;

## Notes

The authors declare the following competing financial interest(s): D.P. is a co-founder of and holds equity in Turia Biosciences.

