## Supporting Information for "Multi-Layer Functional Metabolomics Reveals Chemical Interactions Between the Plant-Associated Bacterium *Serratia marcescens* and *Phytophthora*"

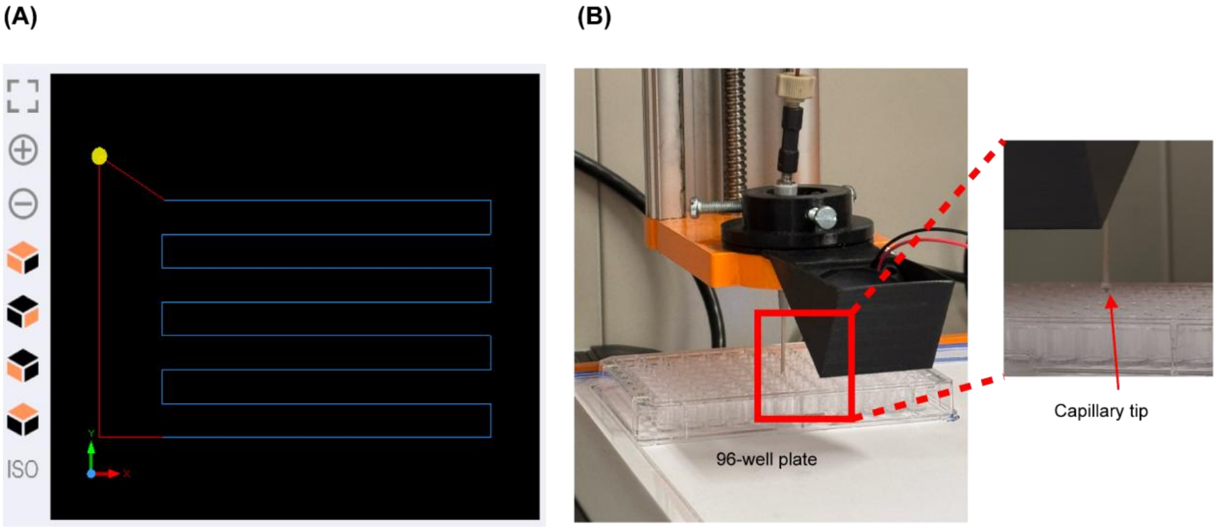

**Figure S1:** Technical details of the microspotter setup. (A) Programmed serpentine ('snake') spotting path used for sequential deposition across the 96-well plate. (B) Photograph of the microspotter during fraction deposition into a 96-well plate, with inset showing a close-up of the capillary tip during spotting.

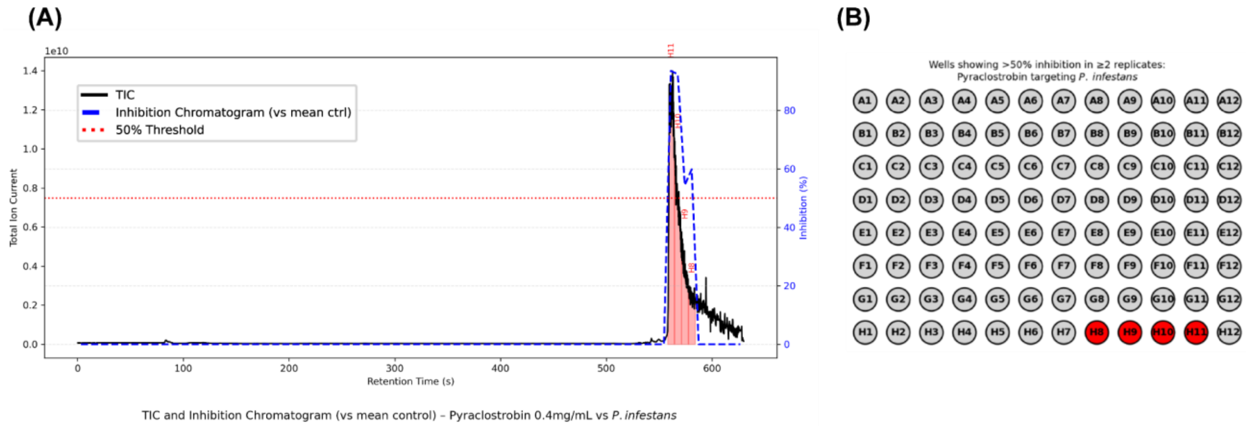

**Figure S2:** Validation of the microfractionation–bioassay workflow using pyraclostrobin as a positive control. (A) Total ion chromatogram (TIC) overlaid with the reconstructed inhibition profile after microfractionation of pyraclostrobin. The red dashed line indicates the 50% inhibition threshold. (B) Map of wells showing ≥50% inhibition across two replicate plates; red wells indicate positive responses.

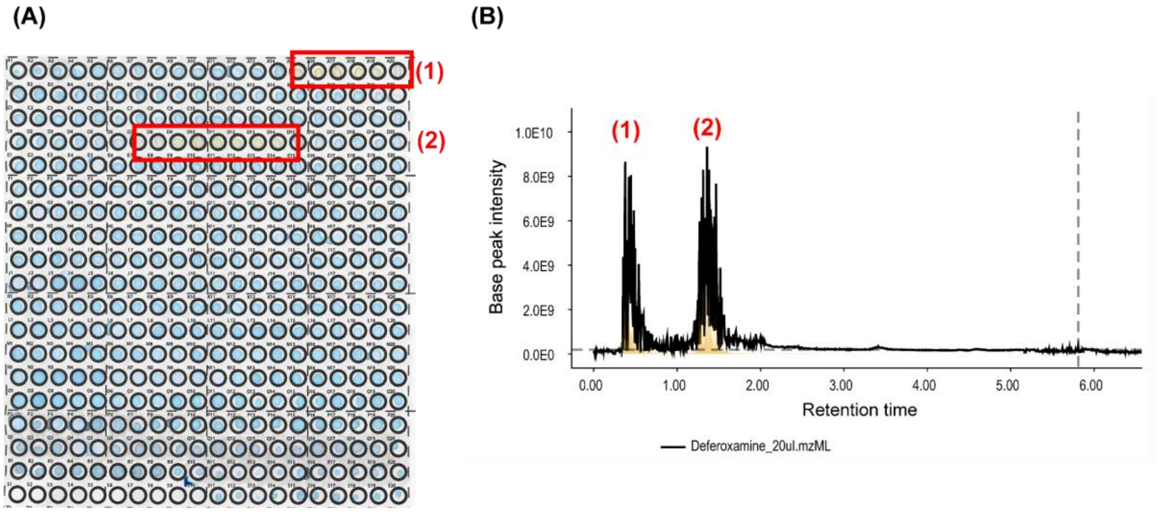

**Figure S3:** Validation of the microfractionation–CAS workflow using deferoxamine as a positive control on microfluidic paper-based analytical devices (µPADs), as described by Geibel *et al.*[1] (A) CAS assay of the microfractionated deferoxamine sample, showing two CAS-positive regions, indicated as (1) and (2). (B) Corresponding LC–MS total ion chromatogram (TIC), showing two chromatographic regions aligned with the CAS-positive responses. Both regions contained deferoxamine-related ions, including $[M+H]^+$ at *m*/*z* 561.3599 and $[M+2H]^{2+}$ at *m*/*z* 281.1835.

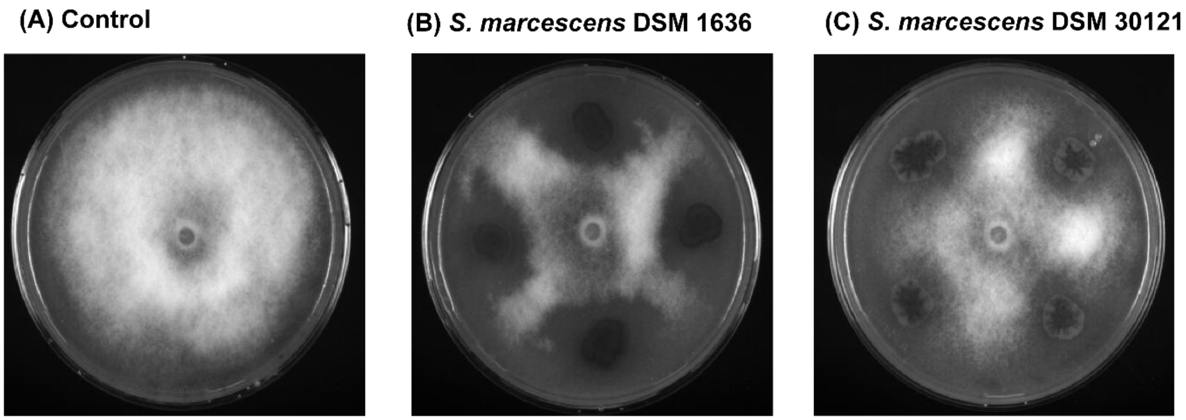

**Figure S4:** Dual-culture antagonism of *S. marcescens* strains against *P. infestans*. (A) Control plate showing unrestricted growth of *P. infestans.* (B) Dual culture of *P. infestans* with *S. marcescens* DSM 1636. (C) Dual culture of *P. infestans* with *S. marcescens* DSM 30121. Reduced pathogen growth in the presence of both bacterial strains indicates antagonistic activity.

**(A) Control** **(B) Intracellular extract** **(C) Extracellular extract**

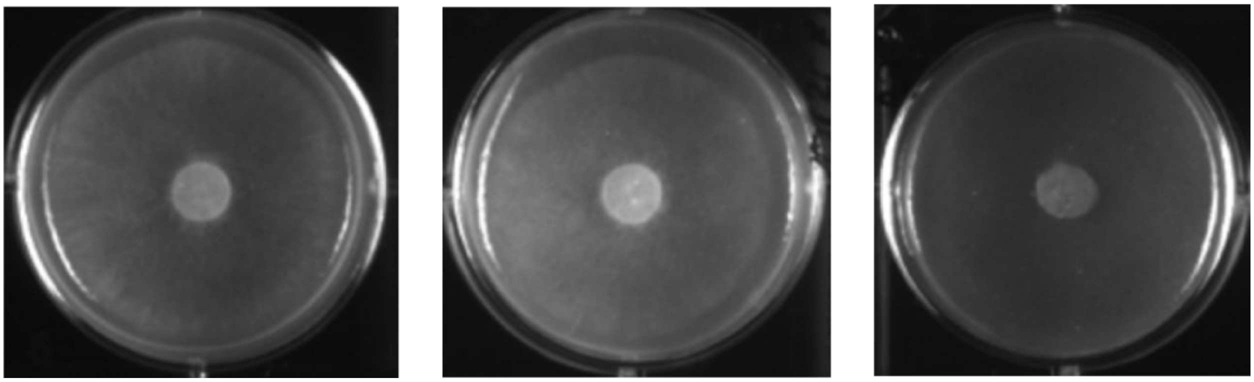

**Figure S5:** Comparison of intracellular and extracellular extracts from *S. marcescens* DSM 30121 against *P. infestans*. (A) Vehicle control containing 0.1% DMSO. (B) *P. infestans* exposed to the intracellular extract at 0.5 mg/mL. (C) *P. infestans* exposed to the extracellular extract at 0.5 mg/mL. The extracellular extract showed a stronger reduction in pathogen growth than the intracellular extract.

**(A)** **(B)**

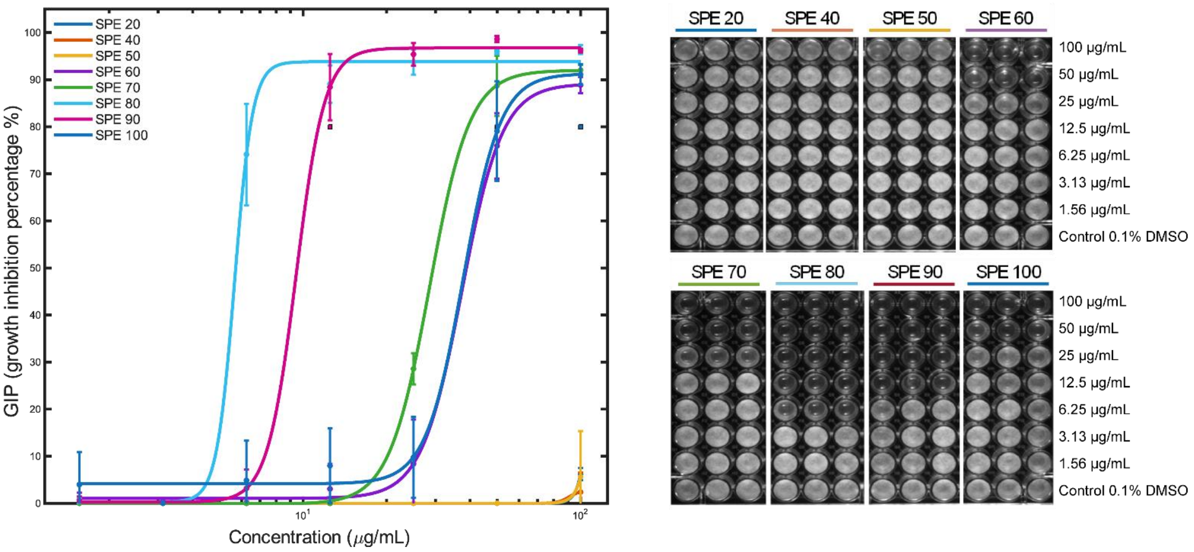

**Figure S6:** Dose–response activity of SPE fractions from *S. marcescens* DSM 30121 against *P. infestans*. (A) Dose–response curves for SPE20–SPE100 fractions across the tested concentration range. Data are shown as mean ± SD. (B) Representative assay plates corresponding to the tested SPE fractions and concentrations; the vehicle control contained 0.1% DMSO.

| SPE fraction | DSM 30121 $IC_{50}$ [µg/mL] | DSM 1636 $IC_{50}$ [µg/mL] |
|---|---|---|
| 20 | n.r. | n.r. |
| 40 | n.r. | n.r. |
| 50 | n.r. | n.r. |
| 60 | 40.6 | 55.6 |
| 70 | 30.8 | 14.1 |
| 80 | 5.6 | 2.6 |
| 90 | 9.9 | 6.0 |
| 100 | 40.1 | 34.9 |

**Table S1:** Calculated $IC_{50}$ values of SPE fractions from *S. marcescens* DSM 30121 and DSM 1636 against *P. infestans*. $IC_{50}$ values are expressed in µg/mL and were obtained from four-parameter logistic dose–response fits. “n.r.” indicates that 50% inhibition was not reached within the tested concentration range.

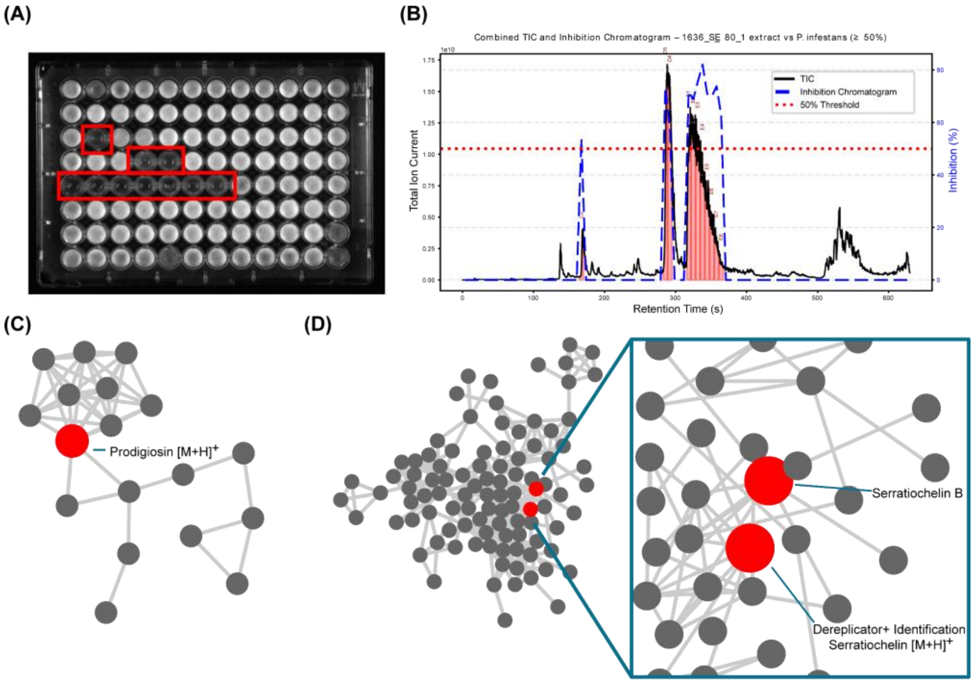

**Figure S7:** Microfractionation-resolved bioactivity profiling and molecular networking of the 80% MeOH SPE fraction from *S. marcescens* DSM 1636 against *P. infestans*. (A) Bioassay plate after microfractionation; red boxes indicate wells showing ≥50% growth inhibition. (B) Total ion chromatogram (TIC) overlaid with the reconstructed inhibition profile; the red dashed line indicates the 50% inhibition threshold. (C) Molecular-network cluster containing prodigiosin, annotated in GNPS2. Prodigiosin corresponded to the most abundant feature across active wells E1–E8. (D) Serratiochelin-related molecular-network cluster. Features matching the retention-time windows associated with activity were prioritized, including serratiochelin A, annotated in GNPS2 and associated with wells D4–D5, and serratiochelin B, assigned based on MS/MS and corresponding to the most abundant feature in well C2.

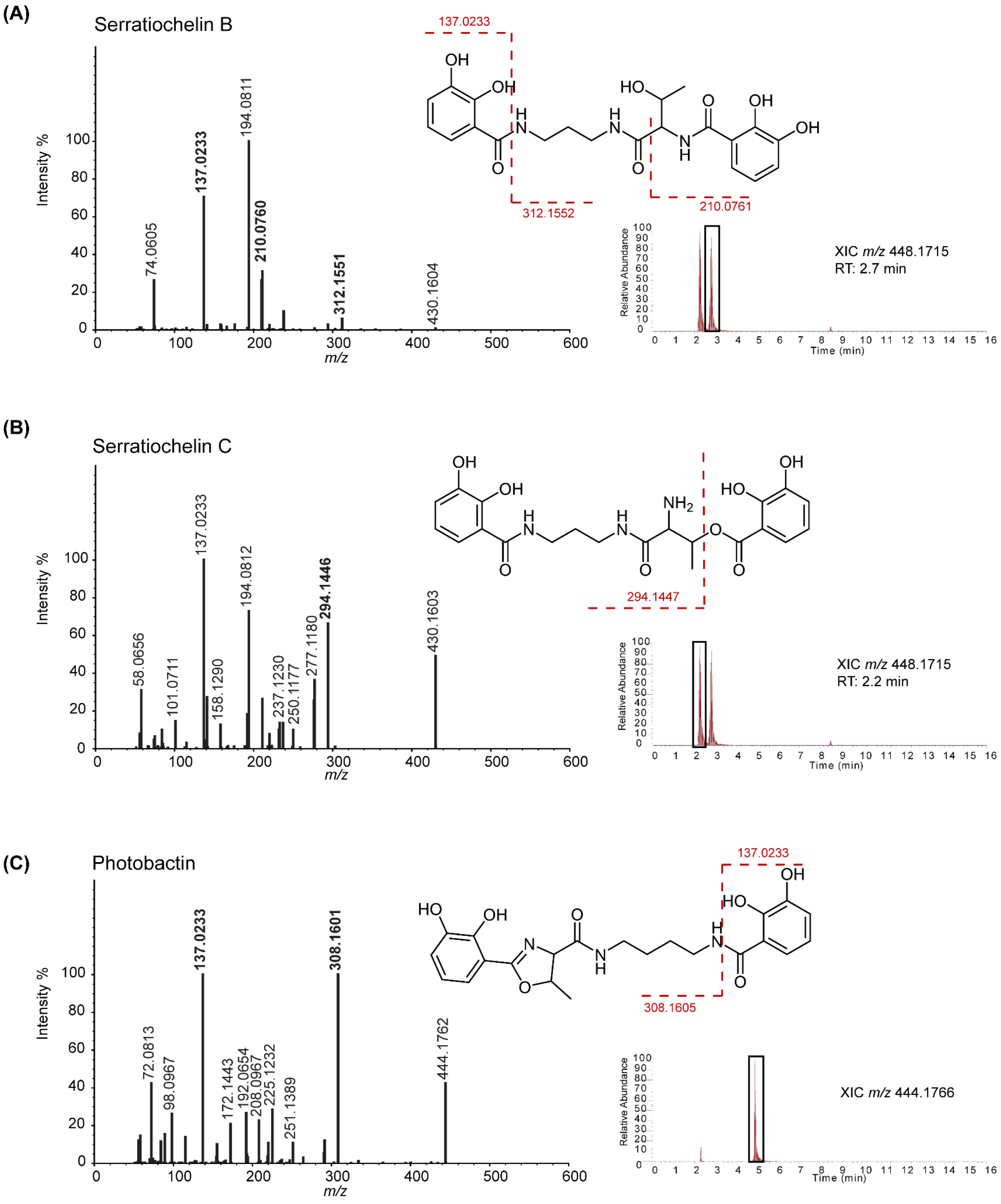

**Figure S8:** MS/MS-based support for the annotation of serratiochelin-related metabolites detected in the 80% MeOH SPE fraction of *S. marcescens* DSM 30121. Positive-ion-mode MS/MS spectra are shown together with the proposed chemical structures and corresponding extracted ion chromatograms. (A) Serratiochelin B (*m/z* 448.1715 RT: 2.7). Selected theoretical fragment ions derived from the proposed structure are indicated in red at *m/z* 137.0233, 210.0761, and 312.1552. The corresponding experimentally observed fragment ions occurred at *m/z* 137.0233, 210.0760, and 312.1551, corresponding to mass errors of 0.00, −0.48, and −0.32 ppm, respectively. (B) Serratiochelin C (*m/z* 448.1715 RT: 2.2). The proposed theoretical fragment at *m/z* 294.1447 corresponded to an experimentally observed fragment at *m/z* 294.1446, with a mass error of −0.34 ppm. (C) Photobactin (*m/z* 444.1766). Proposed theoretical fragment ions at *m/z* 137.0233 and 308.1605 corresponded to experimentally observed ions at *m/z* 137.0233 and 308.1601, with mass errors of 0.00 and −1.30 ppm, respectively. Red dashed lines indicate the proposed structural origins of the selected theoretical fragment ions, while labeled peaks in the MS/MS spectra represent experimentally observed fragment ions. The extracted ion chromatograms (XICs) show the chromatographic profiles of the respective precursor ions.

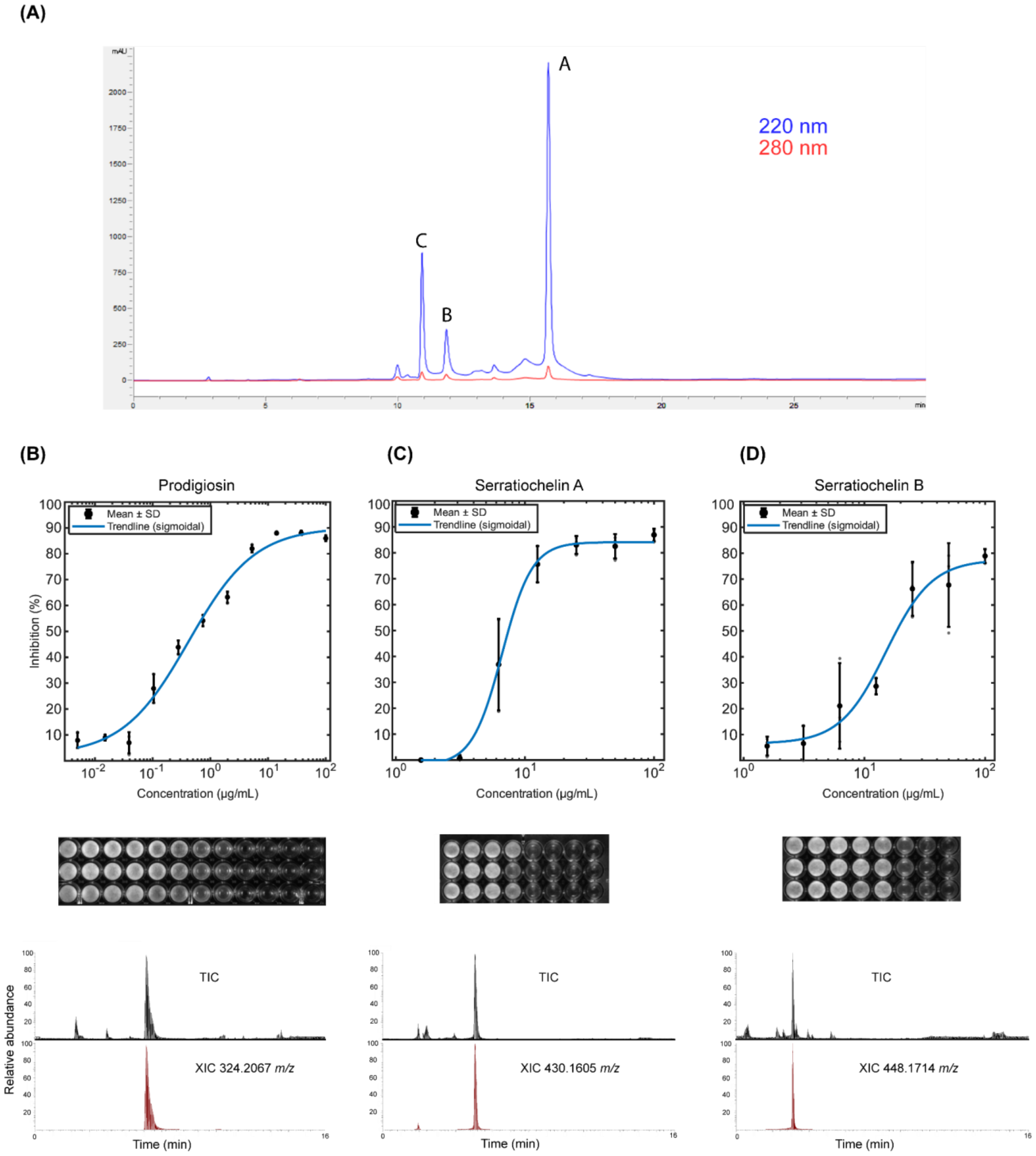

**Figure S9: Bioactivity validation of prodigiosin and purified serratiochelins against *P. infestans*.** (A) HPLC chromatograms of the 80% MeOH SPE fraction of *S. marcescens* DSM 30121 recorded at 220 nm (blue) and 280 nm (red). Peaks assigned to serratiochelins C, B, and A eluted at 10.9, 11.9, and 15.8 min, respectively. (B–D) Bioactivity and LC–MS profiles of the commercial prodigiosin standard (B), purified serratiochelin A (C), and purified serratiochelin B (D). Serratiochelins were purified from the extracellular extract of *S. marcescens* DSM 30121.

For each compound, the upper panel shows the concentration–response curve, with growth inhibition expressed as mean ± SD and fitted using a four-parameter logistic model. The middle panel shows representative assay wells across the tested concentration range. The lower panel shows the total ion chromatogram (TIC) and extracted ion chromatogram (XIC) for the corresponding target ion: *m/z* 324.2067 for prodigiosin, 430.1605 for serratiochelin A, and 448.1714 for serratiochelin B. Estimated $IC_{50}$ values were 0.5, 7.4, and 18.5 µg/mL, respectively.

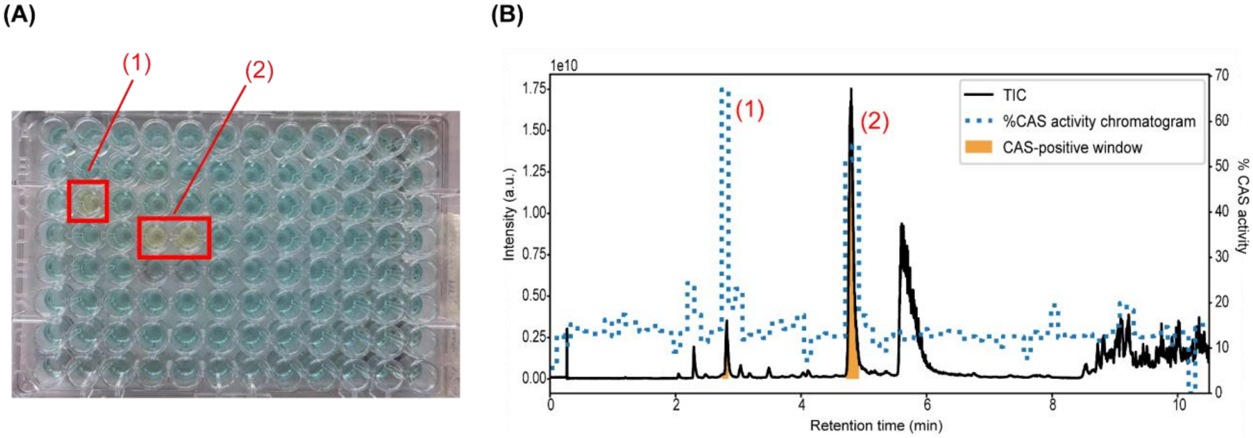

**Figure S10:** Microfractionation–CAS profile of *S. marcescens* DSM 1636. (A) CAS assay plate after LC microfractionation, showing two CAS-positive regions indicated as (1) and (2). (B) Total ion chromatogram (TIC) overlaid with the reconstructed CAS-activity profile. Region (1) corresponded to serratiochelin B ($[M+H]^+$, *m/z* 448.1714), whereas region (2) corresponded to serratiochelin A ($[M+H]^+$, *m/z* 430.1612). Both features showed CAS activity consistent with iron-chelating behavior. Prodigiosin did not produce a positive CAS response under the assay conditions used.

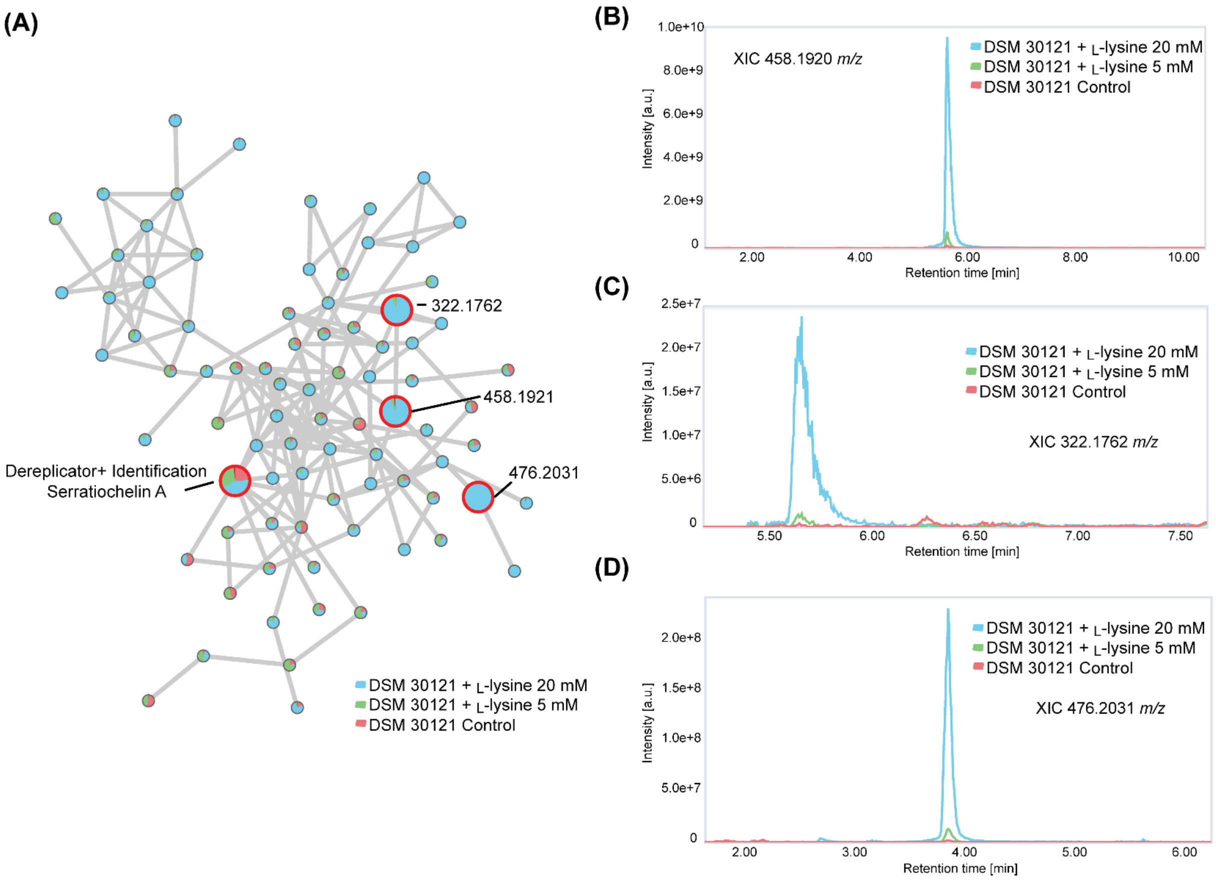

**Figure S11:** L-Lysine-dependent enrichment of serratiochelin-related features in *S. marcescens* DSM 30121. (A) Feature-based molecular network showing the relative abundance of metabolites detected in cultures supplemented with 20 mM L-lysine (blue), 5 mM L-lysine (green), or without added L-lysine (red). Pie charts represent the relative abundance of each feature across the three conditions. Numerous features within the molecular network showed increased relative abundance following L-lysine supplementation, particularly at 20 mM, indicating that the response extended beyond a single metabolite. Selected nodes are highlighted in red, including Dereplicator+-annotated serratiochelin A, serratiochelin G (*m/z* 458.1921), and serratiochelin-related features at *m/z* 322.1762 and 476.2031. (B–D) Extracted ion chromatograms of serratiochelin G, *m/z* 322.1762, and *m/z* 476.2031, respectively, showing increased signal following L-lysine supplementation, with the strongest enrichment observed at 20 mM L-lysine.

### Supplementary methods

#### Dual-culture Antagonism Assay:

A 0.5 cm diameter mycelial plug from a 5-day-old *P. infestans* culture was placed in the center of a V8 agar plate and incubated at 17 °C for 48 h. Subsequently, 5 µL of a *S. marcescens* DSM 1636 or DSM 30121 cell suspension ($1 \times 10^5$ CFU/mL) was inoculated at four equidistant positions located 3 cm from the center of the *P. infestans* plug. Plates were then incubated at 17 °C for an additional 6 days. Each treatment was performed with at least three biological replicates.

#### General LC–MS/MS Analysis

Analyses were performed using a Vanquish UHPLC system coupled to a Q Exactive HF quadrupole-Orbitrap mass spectrometer equipped with a heated electrospray ionization (HESI) source (Thermo Fisher Scientific, Bremen, Germany). Chromatographic separation was achieved on a Kinetex EVO C18 column (50 × 2.1 mm, 1.7 µm, 100 Å; Phenomenex, Torrance, USA), maintained at 25 °C, with the autosampler at 14 °C. Mobile phases consisted of water (A) and acetonitrile (B), both LC–MS grade and containing 0.1% formic acid. The flow rate was 0.5 mL $min^{-1}$, and the injection volume was 5 µL. The gradient increased linearly from 5% to 50% B over 8 min and to 99% B at 10 min, followed by a hold until 13 min, a return to 5% B at 13.1 min, and re-equilibration until 16 min. Data was acquired in positive-ion mode using data-dependent acquisition. Full MS scans covered *m/z* 150–1500 at a resolution of 30,000, followed by MS/MS acquisition of the five most intense precursor ions at a resolution of 15,000, using stepped normalized collision energies of 20, 25, and 30 eV. Source settings were 3.5 kV spray voltage, 250 °C capillary temperature, 400 °C probe heater temperature, sheath gas 50, auxiliary gas 12, sweep gas 1, and S-lens RF level 50.

#### Mass Spectral Data Processing Using mzmine for FBMN:

LC–MS/MS mzML files were processed using mzmine (version 4.5.37) following a complete feature-based molecular networking (FBMN) workflow.

- $MS^1$ and $MS^2$ mass detection were performed using noise thresholds of $3 \times 10^5$ and $1 \times 10^4$, respectively.

- Chromatograms were constructed with the ADAP chromatogram builder using a minimum group size of 5 scans, a group intensity threshold of 2 × $10^5$, a minimum absolute height of 1 × $10^6$, and an *m/z* tolerance of 10 ppm (0.002 *m/z*).
- Peak deconvolution was carried out with the Minimum Search resolver, applying a chromatographic threshold of 85%, peak duration limits of 0.01–2.0 min, a minimum of 5 scans per peak, a minimum absolute height of 1 × $10^6$, a peak top/edge ratio of 1.2, and MS/MS scan pairing tolerances of 10 ppm in *m/z* and 0.2 min in RT.
- Isotopic features were annotated using the isotope grouper (5 ppm *m/z* tolerance, 0.01 min RT tolerance, maximum charge 2, monotonic shape required, representative isotope = most intense) together with the isotope finder module (elements: H, C, N, O, S, Cl, Br; 10 ppm tolerance).
- Feature alignment across all samples was performed with the Join Aligner using an *m/z* tolerance of 10 ppm, an RT tolerance of 0.15 min, and weights of 3 (*m/z*) and 1 (RT), without requiring identical charge or feature IDs.
- Rows corresponding to $^{13}C$ isotopes were filtered out using a 3 ppm tolerance. Missing values were recovered through gap filling (Peak Finder) with an intensity tolerance of 20%, an *m/z* tolerance of 5 ppm, and an RT tolerance of 0.05 min.
- Duplicate features were resolved using the Duplicate Filter with tolerances of 3 ppm (*m/z*) and 0.1 min (RT), using the NEW_AVERAGE merge mode.
- For molecular networking, features were exported using the GNPS FBMN export module, merging MS/MS spectra across samples and applying an *m/z* deviation of 5 ppm, a cosine similarity threshold of 0.7, feature intensity based on peak area, and simple CSV formatting. Output files (.mgf and .csv) were used for FBMN in GNPS2.

**GNPS2 FBMN Workflow Parameters:**

Feature-based molecular networking (FBMN) was performed on GNPS2 using the feature table (.csv) and spectral file (.mgf) exported from mzmine. The analysis used the feature_based_molecular_networking_workflow with the following parameters. Precursor and fragment ion mass tolerances were set to 0.01 Da. Network edge creation required a minimum cosine similarity of 0.7 and at least 5 matched fragment peaks. For library identification, the same cosine threshold (0.7) and minimum matched peaks (5) were applied, selecting only the highest-scoring library hit per feature. Analog search was disabled (library_analog_search = 0), and the maximum allowed precursor shift in both molecular networking and library components was 1999 Da, the default GNPS setting. Spectral filtering options included precursor filtering (precursor_filter = 1) and a window filter width of ±50 Da (window_filter = 1), retaining the top 10 fragments per 50 Da bin (topology_topk = 10). The maximum network component size was set to 100 nodes to avoid oversized clusters. No normalization was applied to feature intensities (normalization = None).

**HPLC Purification of Serratiochelins:**

The 80% MeOH SPE fraction of *S. marcescens* DSM 30121 was subjected to HPLC separation using a Waters system equipped with a 1525 binary pump with an integrated degasser, a Rheodyne 7725i injection valve, and a 996 photodiode array detector. Separation was performed on a Phenomenex Luna® C18(2) column (250 × 4.6 mm, 5 μm) at a flow rate of 0.5 mL/min. Mobile phases consisted of acetonitrile containing 0.05% formic acid (A) and water containing 0.05% formic acid (B). The gradient was programmed as follows: 0–2 min, 10% A; 2–18 min, 10–30% A; 18–25 min, 30–100% A; and 25–30 min, 100% A. The column was then returned to the initial conditions and re-equilibrated for 5 min. UV absorbance was monitored at 220 and 280 nm.

**LC–MS/MS Processing for Ion Identity Molecular Networking (IIMN):**

LC–MS/MS mzML files from Fe(III) infusion experiments were processed in mzmine (version 4.5.37) following the recommended workflow for Ion Identity Molecular Networking (IIMN). $MS^1$ and $MS^2$ mass detection were performed using noise thresholds of $3 \times 10^5$ and $1 \times 10^4$, respectively. Chromatograms were generated using the ADAP chromatogram builder with a minimum group size of 5 scans, group intensity threshold $2 \times 10^5$, minimum absolute height $1 \times 10^6$, and an *m/z* tolerance of 10 ppm. Peak deconvolution was carried out using the Minimum Search resolver, applying a chromatographic threshold of 85%, peak duration limits of 0.01–2.0 min, a minimum of 5 scans per peak, and a minimum height of $1 \times 10^6$. Isotopic features were grouped using the isotope grouper (5 ppm *m/z* tolerance; 0.01 min RT tolerance; max charge 2; monotonic shape required). Feature alignment was performed using the Join Aligner with a tolerance of 10 ppm (*m/z*) and 0.15 min (RT). Gap filling used the Peak Finder module with an intensity tolerance of 20%, *m/z* tolerance 5 ppm, and RT tolerance 0.05 min. Duplicate features were removed using a 3 ppm *m/z* and 0.1 min RT tolerance. For IIMN, ion identity relationships were generated using the Ion Identity Networking module. The primary ion pair was

$[M+H]^+/[M+Fe-2H]^+$ ($\Delta m/z$ = +52.9115). Optional secondary pairs, where present, were $[M+Na]^+/[M+Fe-2H]^+$ and $[M+K]^+/[M+Fe-2H]^+$.

The following ion-identity settings were applied:

- *m/z* tolerance for ion identity pairing: 5 ppm
- RT tolerance: 0.2 min
- Maximum charge: +2
- Require $MS^2$ for both nodes: false
- Minimum height for paired features: $1 \times 10^6$
- Ion identity intensity correlation threshold: 0.7

IIMN-enabled feature tables and MS/MS summary files were exported using the GNPS IIMN export module, merging spectra across samples and producing the required .mgf, .csv, and ion-identity edges for molecular networking in GNPS2. Networks were visualized in Cytoscape.
